# Students Attitudes Surrounding STEM: A Social Cognitive Career Theory Instrument for High School

**DOI:** 10.1101/2021.11.29.470294

**Authors:** EmilyKate McDonough, Kayle S. Sawyer, Jessica Wilks, Berri Jacque

## Abstract

To broaden participation in science, technology, engineering, and math (STEM), we must understand the factors that shape perspectives and beliefs around career selection. Good measurement of these factors is crucial to quantify how effectively educational interventions impact student attitudes towards STEM. Adolescents are particularly suited for quantifying intervention efficacy because students build their identities during these formative years and make important career choices. To better quantify intervention efficacy at the high school level, we developed an instrument entitled Student Attitudes Surrounding STEM (SASS), which builds upon the social cognitive career theory (SCCT) framework for understanding career selection. Questionnaire responses were collected from 932 high school students and split into samples of 400 for exploratory factor analysis and 532 for confirmatory factor analysis. The 37 questions clustered into six factors: Self-Efficacy-Experience, Self-Efficacy-Academic, Outcome Expectations, Interests, Negative Perceptions of Scientists, and Career Awareness. Adequate construct validity for the factors indicated in the SASS model was suggested by the fit indices and theoretical considerations. Furthermore, the analyses supported criterion validity, internal consistency, and test-retest reliability. This tool represents a novel integration of three latent variables into SCCT: Negative Perceptions of Scientists, Career Awareness, and an experience factor for Self-Efficacy.

## Introduction

Educating students in science, technology, engineering, and math (STEM) is essential for their development into informed citizens and active members of the 21st century workforce. Additionally, the skills used in STEM help people lead fulfilling lives, make sensible personal decisions, and support productive civic and community engagement. However, there is growing concern of a ‘leaky STEM pipeline,’ referring to the high rate of attrition from STEM majors and graduate degrees. Over the past couple of decades in the United States, 40%-50% of undergraduate students who enrolled in STEM majors left the field before their scheduled graduation, or shortly thereafter (Chen, 2013; PCAST, 2012). Particularly high rates of attrition have been seen in the populations already underrepresented in STEM, such as women, people of color, and first generation college students (Anderson & Kim, 2006; Gayles & Ampaw, 2014; Hill et al., 2010; Shaw & Barbuti, 2010). This loss of talent within the STEM workforce leads to the reduction of diverse perspectives and ideas, and limited representation of community stakeholders that are essential for progress in science and medicine.

Just as the paths people take to obtain a STEM career are varied, so too are their reasons for exiting the STEM pipeline. Numerous contextual factors, such as access to role models, peer support, institutional environment, and discrimination contribute to a person’s decision to leave STEM (Chang et al., 2011; Fouad et al., 2010; Gayles & Ampaw, 2014; Griffith, 2010; Hill et al., 2010; Wang, 2013). Additionally, various individual factors impact which students are likely to leak out of the STEM pipeline. For example, undergraduate students majoring in STEM fields tend to switch to a non-STEM major if they have lower confidence in their STEM abilities, and this is especially true for women and first generation college students (Shaw & Barbuti, 2010; Wang, 2013). Another factor linked to withdrawal from STEM comes from a waning interest in the field, due to a lack of knowledge surrounding potential STEM careers (Cohen et al., 2013). If students don’t know what careers are available, or what a STEM career actually entails, they may be less likely to pursue one.

While the leaky pipeline term has been criticized for its suggestion that there is a single trajectory to a STEM career (Cannady et al., 2014), the imagery evoked by the term has increased awareness of the attrition problem and prompted the development of numerous STEM education interventions worldwide (UMass Donahue Institute, 2011; van den Hurk et al., 2019). Interventions that effectively retain students in STEM build knowledge and influence student trajectories by fostering healthy self-perceptions and beliefs. To quantify the success of these educational interventions, instruments measure characteristics such as domain knowledge, extracurricular knowledge, aspects of self-perception, and aspirations. Data from these instruments can identify which student groups benefit from the intervention, allowing an educational program to be implemented for those with the greatest need and for those who would receive the greatest benefit. Finally, ideal metrics reveal the efficacy of the intervention in “real time”, allowing for iterative curricular improvements.

In summary, to broaden participation in STEM, careful measurement of educational interventions’ impact on student perspectives is essential. Adolescence is a critical period in which to assess interventions, because students form beliefs about themselves and make enduring choices regarding career selection. Therefore, a good instrument must be sensitive to such interventions, by measuring beliefs and attributes that are not static traits, and it must be short enough to make it practical for teachers to implement in a classroom setting. In this manuscript we describe the development and provide validity evidence for a new questionnaire, which we call Student Attitudes Surrounding STEM (SASS). This tool provides researchers and teachers with a short, efficient, and practical means to measure the impact of educational interventions in high school classroom settings.

## Background and Theoretical Framework

### Social Cognitive Career Theory

In 1994, Robert W. Lent, Steven D. Brown, and Gail Hackett proposed the social cognitive career theory (SCCT) as a unified framework through which to understand career choices. The theory is based on the work of Albert Bandura’s social cognitive theory, which examines the relationship between personal factors (e.g., beliefs, attributes), environmental factors (e.g., feedback, culture), and behavior (Bandura, 1986). At its core, SCCT measures the relationships between self-efficacy (“I am good at X”), outcome expectations (“If I do X, this will happen”), interests (“I’m interested in X”), and goals (“I plan to do X”), as described by Lent and colleagues (1994, 2000). For example, a STEM-specific model suggests that someone who considers themselves good at math, and who expects positive outcomes from a math-related career, would be more interested in math, set goals to study math, and persist within STEM. That is, higher confidence and belief in favorable outcomes will help students set goals and persist within STEM. Thus, self-efficacy, outcome expectations, and interests, influence career-related goal-setting and behavior.

While the four social cognitive factors (self-efficacy, outcome expectations, interests, and goals) represent the center of SCCT, they do not exist in isolation. People respond to their environments and incorporate social feedback as they develop beliefs about themselves. Therefore, researchers have examined how additional constructs influence these core factors, and ultimately, career choice. Lent and colleagues (1994, 2000) proposed that personal characteristics (e.g., gender, race/ethnicity, emotional affect, socioeconomic status [SES]), background factors (e.g., family expectations, role model exposure), and learning experiences influence these cognitive factors and mediate their impact on career choice. Additionally, career choice outcomes are influenced by external factors such as the supports (e.g., social network) and barriers (e.g., discrimination) one might expect to receive as a consequence of choosing a STEM career (Lent et al., 1994, 2000). While no single environmental element determines outcomes by itself, each can have a powerful impact. For example, high SES, which allows for greater access to learning experiences and career-support resources, serves as one form of support that influences career choices and persistence. However, people with financial resources do not always succeed in attaining their career goals, whereas people with few resources have succeeded none-the-less. People respond to their environment differently for a wide variety of reasons, as captured by the various environmental inputs of the SCCT framework.

SCCT has been used to examine STEM career choice in a wide range of populations. The framework has been used for research examining outcomes for low income, prospective first generation college students (Garriott et al., 2013), as well as for examining effects of race, ethnicity and gender (Lent et al., 2005; Navarro et al., 2007; Turner et al., 2019). These individual characteristics are commonly referred to as person inputs. While mostly explored at the undergraduate and graduate level, the SCCT model has been validated for various age groups, including high schoolers, college students, and people early in their careers (Fouad & Santana, 2017; Lent et al., 2018). Indeed, the SCCT framework is a powerful tool for understanding career choice among different populations. However, the relationships between the SCCT constructs have varied when examining different populations, especially with respect to self-efficacy and outcome expectations (Fouad & Santana, 2017). These studies illustrate the framework’s utility, while also highlighting the need to validate the SCCT model within specific contexts.

People begin to consider their future professions starting from a young age, often before beginning their formal education. Throughout childhood and adolescence, changes in self-efficacy, outcome expectations, and interests can impact career choice. Adolescence is a crucial time in people’s lives with respect to their career, because it is when people make impactful decisions, such as whether to attend college. Measuring the SCCT constructs in high school students allows us to understand what motivates potential STEM professionals early in their career trajectory. SCCT research within the high school population has focused on career interest as an endpoint measure, rather than career goals or actions (Lent et al., 2018). This may reflect how interests are a more pertinent measure for adolescents than goals, which may not be well-formed in this population. Science self-efficacy has been correlated with science career interest for students in health career-focused extracurricular programs, although this observation may be specific to the selected population of adolescents who show high interest and achievement in STEM (Peterman et al., 2018). However, even as early as middle school, positive experiences in science and math are associated with higher STEM self-efficacy and career interest (Fouad & Santana, 2017). These studies show that researchers can use SCCT to robustly measure a STEM intervention’s impact on future career choices in adolescent populations.

The literature examining the SCCT latent variable relationships during adolescence has mainly focused on single subjects such as math (Garriott et al., 2013; Lim & Chapman, 2013; Lopez et al., 1997; Lopez & Lent, 1992; Schukajlow et al., 2012) or bioinformatics (Kovarik et al., 2013), or on single disciplines such as science (Stake, 2006; Syed et al., 2012), but these narrowly focused instruments are too numerous to review here. To our knowledge, only three instruments have been developed to investigate the SCCT constructs in the context of all-inclusive STEM with high school students: the STEM Career Interest Survey (STEM-CIS; Kier et al., 2014), the Student Attitudes Toward STEM (S-STEM; Unfried et al., 2015), and the Student Interest and Choice in STEM (SIC-STEM; Roller et al., 2020). STEM-CIS was used to assess Taiwanese high school STEM career interest in the context of SCCT, but items were not examined using SCCT constructs: self-efficacy, outcome expectations, interests, goals, supports, and person inputs (Kier et al., 2014). Instead, latent variables containing representative SCCT items were created for each discipline (science, technology, engineering, and math), and validity evidence was examined for the discipline-related factor structure. Likewise, S-STEM, and its derivative SIC-STEM, was developed with latent variables for 21st century skills, science attitudes, math attitudes, and a combined scale for science and technology attitudes (Roller et al., 2020; Unfried et al., 2015). Within each latent variable, items were included to assess self-efficacy, outcome expectations, interests, and career goals. Recently, STEM-CIS was adapted for high school students in China, and the factor structure was validated with items grouped by SCCT construct, rather than STEM discipline (Mau et al., 2019). This demonstrates that the 44-item STEM-CIS instrument could be used for evaluation of the SCCT constructs within a high school population. However, it is too long to be practical in some settings, especially considering that we were additionally interested in measuring perceptions of scientists and career awareness, which the STEM-CIS does not include.

### Perceptions of Scientists

In 1957, Margaret Mead and Rhoda Métraux asked approximately 35,000 high school students across the United States to write essays about their views on scientists. Students responding to the writing prompt often described a scientist’s work as admirable and valuable to society. However, this positive view was paired with negative depictions of a scientist when students were asked to describe themselves or a romantic partner as a scientist. For example, one student described a scientist as having “no other interests” and as someone who “neglects his family” (Mead & Métraux, 1957). Research built upon this work found that several of these false stereotypes have persisted for decades (Chambers, 1983; Dikmenli, 2010; Finson, 2002; Pion & Lipsey, 1981; Schinske et al., 2015; Scholes & Stahl, 2020; Wyer et al., 2010). Furthermore, students can be ignorant about what scientists actually do, believing that all scientists conduct dangerous work and shout “Eureka!” upon successful completion of an experiment (Chambers, 1983; Christidou, 2011; Mead & Métraux, 1957; Scholes & Stahl, 2020). These stereotypes of scientists and science have been observed in students across the globe and in various settings (Christidou, 2011). In summary, research spanning 60 years shows that secondary students hold a confused and unrealistic picture of what it means to be a scientist.

The persistence of negative perceptions of scientists is especially concerning as it dissuades people from pursuing STEM, which in turn contributes to the leaky pipeline. Indeed, stronger aspirations of undergraduate students to pursue a STEM career are associated with more positive perceptions of scientists (Wyer, 2003). Furthermore, science self-efficacy and interests (major influencers in career choice) are interconnected with student-held images of scientists (Christidou, 2011). Negative stereotypes of scientists are seen more prevalently in Asian undergraduate students than in their peers (Schinske et al., 2015), demonstrating a connection between person inputs (in this case, race) and perceptions of scientists. Interventions that paint a realistic picture of what it means to be a scientist show promise in addressing the racial and gender disparities within STEM. For example, Yonas and colleagues (2020), described a “Scientist Spotlight” intervention that uses podcast episodes from a diverse group of scientists to impact students’ perception of scientists and their belonging in science.

Developing good measures of students’ stereotypes of scientists is crucial to assess the impact of educational interventions aimed at changing the perception of scientists. Several methods have been used to accomplish this goal, ranging from prompted essays (e.g., Mead & Métraux, 1957) and drawings (e.g., Chambers, 1983) to Likert scale rated questionnaires (Krajkovich & Smith, 1982; Wyer et al., 2010). The questionnaires developed by Krajkovich and Smith (1982) and Wyer and colleagues (2010) included items inspired by imagery and beliefs detailed in the initial work of Mead and Métraux (1957). Both scales included positive (e.g., intelligent and careful) and negative (e.g., unhappy family and limited social life) stereotypes of scientists. While the survey developed by Krajkovich and Smith (1982) was initially specified with a single factor, later work highlighted the independence of negative perceptions (Marshall et al., 2010). These instruments help to uncover the breadth of stereotypes students have about scientists, which can be useful in designing interventions that target specific beliefs. However, teachers were not willing to commit the time required for these instruments in addition to the time needed for our SCCT questions, especially if the information we needed could be obtained more quickly. Accordingly, we decided to build a scale that assesses these perceptions with just a handful of questions, which could be integrated into our SCCT instrument.

### Career Awareness

It is hard to develop an interest in a career if one is unfamiliar with it. While most students are familiar with STEM careers such as doctor or computer programmer, fewer are aware of more specialized jobs such as animal care technician or clinical trial recruiter. Within the high school population, higher STEM career awareness, along with higher STEM self-efficacy, is correlated with stronger desires to pursue a STEM career, demonstrating an interplay between SCCT and career awareness (Blotnicky et al., 2018; Zhang & Barnett, 2015). Furthermore, a study of students from a primarily white, middle class high school showed that adolescents are unaware of the skills and knowledge required for entry into their chosen fields of interest (Johnson, 2000). Assuming that career awareness is associated with privilege, then under-resourced and minority groups would have even less awareness than the white, middle class students. This research highlights the importance of incorporating career-specific knowledge into curricular interventions aimed at increasing interest in STEM careers.

Career knowledge can be quantitatively assessed by directly measuring one’s knowledge of required skills, degrees, or duties for specific careers. For example, the Cognitive Vocational Maturity Test (CVMT) is a highly comprehensive, 120-item, six-construct instrument that assesses career knowledge in middle and high school students. While the CVMT provides a wealth of information about an individual’s career knowledge, the full instrument takes two 45-minute class periods to complete, and is therefore impractical for most settings (Westbrook et al., 1996). Alternatively, some instruments measure specific knowledge about careers, such as what degree or classes are required to enter particular fields (e.g., Blotnicky et al., 2018). Another approach to understanding career awareness and career-related knowledge is to rely on self-reported learning. For example, Salonen and colleagues (2018) used three questions to assess perceived learning following a career awareness intervention: *“I gained knowledge about careers that are new to me,” “This unit helped me understand the responsibility of the described careers,”* and *“This unit helped me to understand what skills are needed in the described careers.”* These measures of perceived learning capture students’ confidence in their career knowledge, which, like self-efficacy, could influence career choices.

In the context of a concise SCCT assessment in high school classrooms, these career awareness scales were not suitable for our needs. The CVMT is too long and is not STEM specific. The scale developed by Blotnicky and colleagues (2018) is reasonably short, but assesses knowledge of education requirements, not knowledge about the careers themselves. Finally, the scale developed by Salonen and colleagues (2018) doesn’t focus on specific careers, and refers to the administration of a specific intervention. Instead, we wanted an efficient instrument that simply asked students to report how much they think they know about a sampling of specific careers.

Consequently, we could assess student breadth of knowledge (and confidence in that knowledge) in a targeted and practical way.

### Rationale for a New Instrument

In this manuscript, we describe a new measurement instrument, SASS, to assess high school STEM career attitudes. Our instrument builds on the SCCT framework. As described above, previous instruments focused on self-efficacy, outcome expectations, interests, and goals, and did not include items to measure career awareness or perceptions of scientists. One instrument, designed to measure the impact of career-focused high school bioinformatics curricula, includes career awareness with the SCCT constructs of self-efficacy and interests (Kovarik et al., 2013). However, this instrument focused on bioinformatics alone and did not include outcome expectations, a central factor in SCCT. We aimed to develop an instrument that integrates the measures of SCCT with career awareness and perceptions of scientists so we could efficiently measure the impact of our classroom intervention on STEM career awareness and perceptions.

In summary, the justification for developing this instrument was twofold. First, the existing perceptions of scientists and career awareness instruments were too long. Administering multiple lengthy surveys is problematic from the perspective of teachers, who have limited time to donate to research studies, and from the perspective of high school students, who may quickly develop survey fatigue, which would threaten the veracity of their responses. Further, if surveys are given in multiple sessions, matching de-identified responses by student can be difficult and result in unmatchable records. Second, we know of no career awareness questionnaire that assesses broad knowledge of specific STEM careers. In this paper, we detail the development and psychometric validation of the use of this new instrument. To our knowledge, this is the first instrument that has integrated career awareness and perceptions of scientists so that relationships of these scales with SCCT scales can be conveniently assessed and analyzed, especially with respect to educational interventions.

## Methods

### Instrument Development

We developed the SASS instrument over the course of two years. Items for SASS were drawn from the SCCT instrument for engineering majors (kindly provided upon request by R.W. Lent; Lent et al., 2005), which included the following latent variables: self-efficacy, outcome expectations, interests, goals, social supports, and social barriers. We adapted items from these latent variables into high school appropriate language. Since we were developing a general STEM version of the SCCT questionnaire, the items were also adjusted to reflect STEM issues more broadly (Table 1 and Table S1). Throughout this manuscript, we use the following capitalized terms to refer to the latent variables in the SASS instrument: Self-Efficacy, Outcome Expectations, Interests, Career Awareness, and Negative Perceptions of Scientists.

**Table 1.**
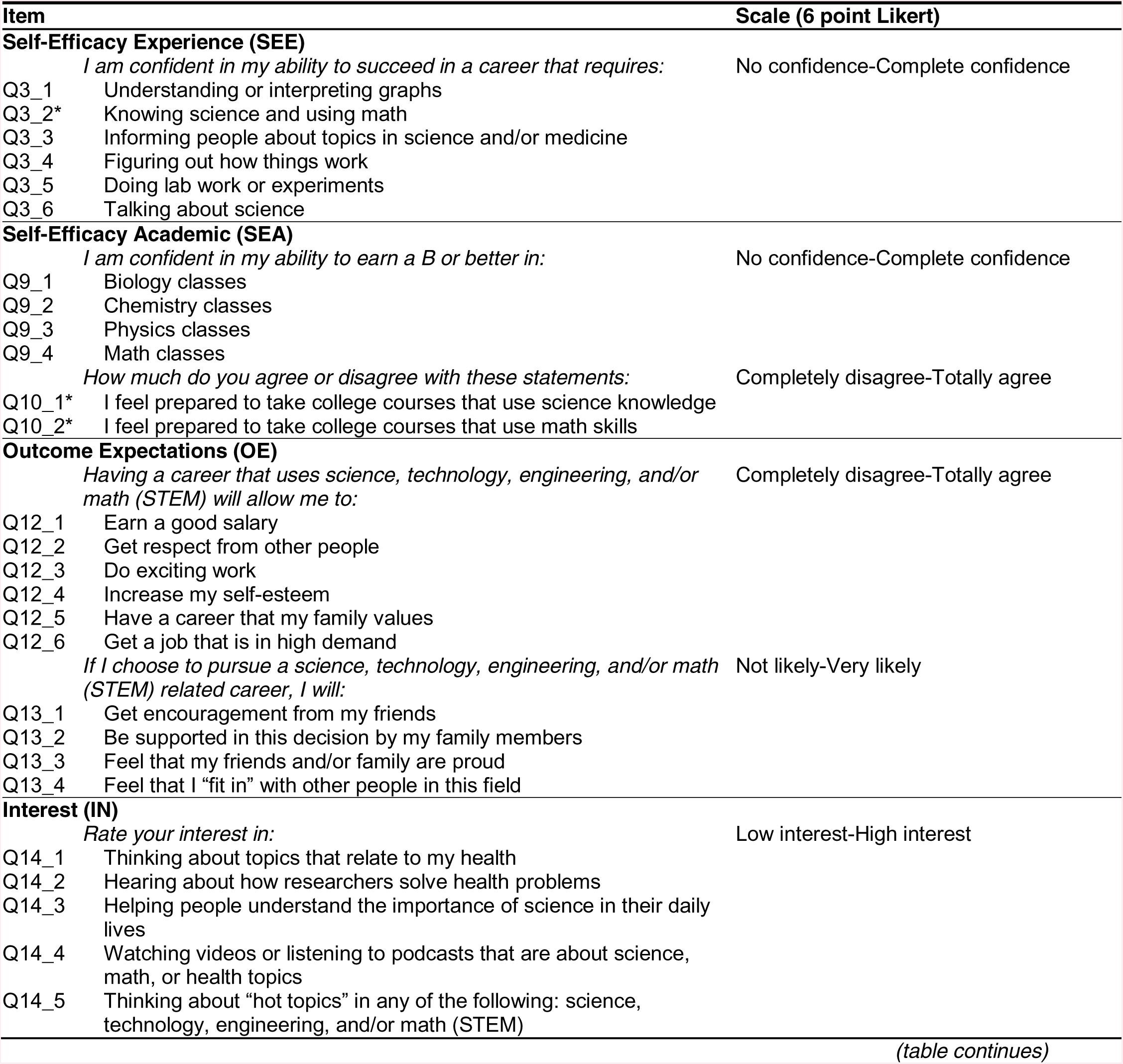
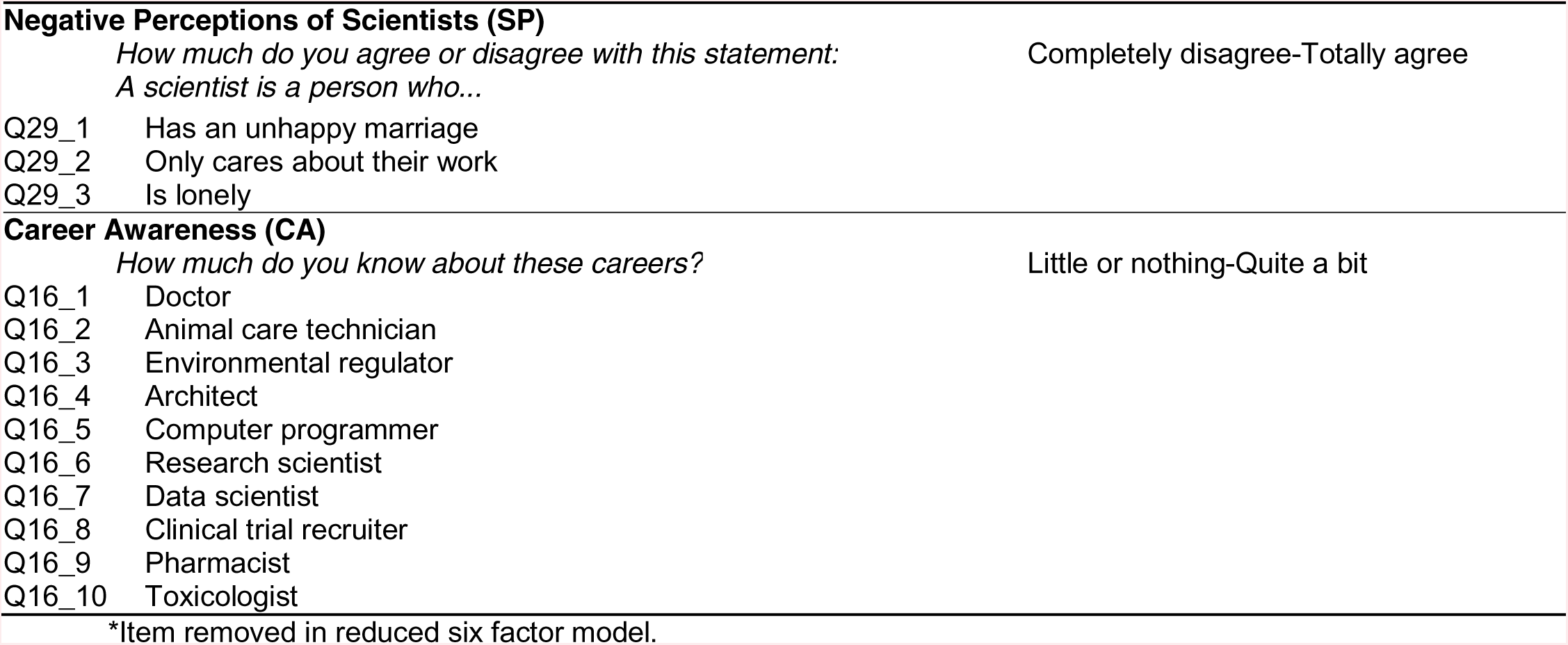
SASS instrument item wording

We developed item content and wording for SASS using an iterative procedure, encompassing four pilot versions with over 1,000 responses. Items were reworded, dropped, or added based on expert opinions, literature and theory, and item performance in exploratory factor analysis (EFA). Following factor analysis, items were considered for rewording or dropping based on three main considerations: 1) coefficients below 0.3 on any expected factor, 2) cross-loading of the item on multiple factors, and 3) loading of the item on an unexpected factor, in contradiction to the a priori model (Costello & Osborne, 2005). During development, items related to the SCCT constructs of social barriers and goals were dropped. Items derived from the social supports variable were strongly correlated with outcome expectation items, which we expected based upon item wording and theoretical considerations. Items from both Outcome Expectations and social supports assessed assumptions about the respect and encouragement the students believed they would receive as a result of choosing a STEM career. For example, two of the items considered as Outcome Expectations are “get respect from other people” and “have a career that my family values,” while two of the items from social supports also refer to judgment by other people: “get encouragement from my friends” and “be supported in this decision by my family members.” Table S1 demonstrates the evolution of several questions from the engineering SCCT (Lent et al., 2005) to our final instrument.

The 40 items in the version of the SASS questionnaire we administered are listed in Table 1. All questions were grouped by predicted construct, rather than randomized, in order to maintain the engineering SCCT survey layout (Lent et al., 2005). Items in Table 1 are listed in the order they were presented to students. The original engineering SCCT was scored on a 10-point or 5-point Likert scale depending on the item. SASS is scored on a 6-point Likert scale for all items. In addition to the SCCT items, we included several demographic (e.g., race, gender, year in school, and school name) and background (e.g., grades in school, intention to go to college, intention to pursue a STEM career, science courses taken, and science activities outside of school) questions in the survey (Table 2 and Table S2). These questions were collected to help understand the environmental factors impacting students’ self-efficacy, outcome expectations, and interests in future studies (as discussed in: Lent et al., 2000).

**Table 2.** Demographics summary statistics

| Variable | Level | EFA (*n* = 400) |  | CFA (*n* = 532) |  |
| --- | --- | --- | --- | --- | --- |
|  |  | *n* | Percent | *n* | Percent |
| Grade | 9th | 78 | 19.5 | 85 | 16.0 |
|  | 10th | 89 | 22.3 | 159 | 29.9 |
|  | 11th | 95 | 23.8 | 129 | 24.2 |
|  | 12th | 136 | 34.0 | 159 | 29.9 |
|  | No response | 2 | 0.5 |  |  |
| Gender | Female | 234 | 58.5 | 269 | 50.6 |
|  | Male | 160 | 40.0 | 248 | 46.6 |
|  | Other / non-binary | 2 | 0.5 | 3 | 0.6 |
|  | Prefer not to answer | 1 | 0.3 | 7 | 1.3 |
|  | No response | 3 | 0.8 | 5 | 0.9 |
| Race | American Indian or Alaskan Native | 4 | 1.0 | 3 | 0.6 |
|  | Asian | 82 | 20.5 | 74 | 13.9 |
|  | Black or African American | 55 | 13.8 | 79 | 14.8 |
|  | Native Hawaiian or Other Pacific Islander | 2 | 0.5 | 4 | 0.8 |
|  | White | 181 | 45.3 | 257 | 48.3 |
|  | Mixed race | 16 | 4.0 | 31 | 5.8 |
|  | Prefer not to answer | 50 | 12.5 | 71 | 13.3 |
|  | No response | 10 | 2.5 | 13 | 2.4 |
| Ethnicity | Not Hispanic / Latino | 291 | 72.8 | 398 | 74.8 |
|  | Hispanic / Latino | 88 | 22.0 | 111 | 20.9 |
|  | Prefer not to answer | 18 | 4.5 | 18 | 3.4 |
|  | No response | 3 | 0.8 | 5 | 0.9 |

The engineering SCCT self-efficacy items refer to confidence about future academic performance (e.g., ability to obtain a B or better in engineering classes) and confidence in the ability to cope with social barriers (e.g., ability to continue in the engineering major without support from professors or not feeling well-liked). We adapted the academic performance items to be appropriate for the high school audience, but we dropped the questions about barrier coping. The retained items reflect the original “self-efficacy for academic milestones’’ questions (Lent et al., 1986). The items related to barrier coping (added subsequently, Lent et al., 2001) did not perform well, perhaps because high school students perceive those concepts differently than college students. Instead, we included questions about confidence in students’ ability to succeed in a career that requires various STEM skills (e.g., figuring out how things work or talking about science). These new items were hypothesized to be part of the Self-Efficacy construct.

The engineering SCCT outcome expectations items refer to job outcomes such as salary, satisfaction, respect, and support. For the SASS instrument, we retained six of the original ten items in this latent variable. These items mostly remained unchanged, with only minor rewording to be more appropriate for high school students. For example, “earn an attractive salary” was changed to “earn a good salary” in order to simplify the language for students. Additionally, because social support items and outcome expectations were correlated, as noted previously, four social supports items were combined with this factor.

The interests items in the engineering SCCT focused on solving problems, working on projects, or learning. Since not all high school students have had the opportunity to work on STEM projects or solve STEM problems, we updated these items to be more reflective of the generic high school student experiences. The SASS interest items use actions such as thinking, hearing, or watching and examine interest levels in health, research, explaining science, and STEM media.

We developed the Negative Perceptions of Scientists construct with the goal of learning how contemporary high school students perceive scientists. As discussed above, previous research demonstrated a correlation between SCCT constructs (self-efficacy, interests, and goals) and perceptions of scientists, supporting the inclusion of perceptions of scientists within an SCCT framework (Christidou, 2011; Wyer, 2003). The Negative Perceptions of Scientists items in SASS were inspired by the work of Mead and Métraux (1957) and we initially included items representing both positive and negative stereotypes of scientists. In preliminary factor analyses with early versions of SASS, these items factored into separate positive and negative constructs, as was observed previously (Marshall et al., 2010). We found that the items related to positive perceptions were highly associated with items from other constructs and did not clearly form a single factor. To make the latent construct most coherent, we retained the items regarding the negative perceptions of scientists and dropped items related to positive stereotypes.

The STEM career awareness items ask students to self-report their knowledge of specific careers (6-point Likert scale; little or nothing - quite a bit). Careers were chosen based on relatedness to a curriculum we created in parallel to SASS (Schneider et al., 2018, 2019). For example, our curriculum introduces students to the careers of animal care technician, research scientist, data scientist, and toxicologist, all of which are included in the latent variable. During pilot testing, we evaluated different lists of STEM careers, and items within the Career Awareness EFA construct still correlated more highly with each other than with items for other constructs.

### Study Participants and Data Collection

All versions (pilot and final) of SASS were tested on high school students in the northeastern United States who were enrolled in a STEM class. Summary statistics of participants included in the reliability and validity testing for the final version are presented in Table 2. Survey instruments and procedures involving research study participants and data management were approved by the Institutional Review Board of Tufts University School of Medicine (#12205 and #533).

Surveys were administered through an online platform; the final version of SASS was hosted on Qualtrics. Responses for each item were requested from participants, but not required. Teachers were asked to administer the survey in class to increase the likelihood that students would take the survey seriously. To account for this, the survey included the following question: “Where are you taking this survey? a) At school in class, b) At school on my own time, c) At home, or d) Other: please specify.” A small subset of students (*n* = 104; 11.2%) reported taking the survey outside of class.

Socio-economic data for students was determined based on school-level reporting of “percent economically disadvantaged.” This data was collected from New York (https://data.nysed.gov; downloaded on Oct 20, 2020) and Massachusetts (https://profiles.doe.mass.edu; downloaded on Oct 19, 2020) state databases (data was not obtained for students from other states). Downloaded files are available at: https://gitlab.com/emcdonough/sass-validation. In both state databases, students were deemed economically disadvantaged based on enrollment in one or more government assistance programs (e.g., food stamps, foster care, Earned Income Tax Credit, Safety Net Assistance Program, etc.).

### Data Analysis

All data analysis was performed using R v4.2.0 (R Core Team, 2013). The code and data used to generate results for this paper are available at: https://gitlab.com/emcdonough/sass-validation. Participants with any missing data (*n* = 44; 4.4%) were removed from the dataset. Of these, 33 (3.3%) withdrew from the study as indicated by failing to finish the survey and 10 (1.0%) participants were missing responses to up to four items. One participant took over 7 days to complete the survey, and only completed the second half. Two methods, similar to those reported previously (Adams et al., 2006; Semsar et al., 2011), were used to assess whether students provided authentic responses. First, students exhibiting string response behavior (e.g., answering 95% 5s or 1s; *n* = 3) were removed from the dataset. Second, the length of time a student took to fill in the survey was automatically recorded by Qualtrics, and this was used to remove students who likely responded to questions without reading them. Based on data from three researchers, an average of 3 seconds per item was determined as the quickest a respondent could fill out the SASS and associated demographic questions and comprehend what they were reading. Therefore, students who completed the survey more quickly than 198 seconds (*n* = 10) were removed from the dataset. The final sample included 932 high school students.

### Validity Testing

Content validity for all items was assessed by six experts in education research, qualitative and quantitative data analysis, psychometric research, and laboratory science (that is, the authors and consultants). Experts were asked to check items for sufficiently simple language, clarity, and relevance to the high school audience. Additionally, experts reviewed items to ensure they did not have multiple interpretations. Any items chosen for rewording were reevaluated by the expert group during meetings, and rewritten collaboratively.

The internal structure for the survey was identified and confirmed using factor analysis to evaluate construct validity. Due to the addition of our two new latent variables, Negative Perceptions of Scientists and Career Awareness, we decided to first perform an EFA. Following the EFA, the factor structure was confirmed using confirmatory factor analysis (CFA). The sample was randomly split to assign 400 participants to the EFA sample and 532 to the CFA sample. These sample sizes are considered robust for factor analysis (Guadagnoli & Velicer, 1988; Rouquette & Falissard, 2011).

Each sample was first tested for factorability using three methods as recommended by other authors (Field, 2013; Knekta et al., 2019). First, visual examination of an inter-item correlation plot (generated using the “corrplot” package for R, version 0.84; Wei & Simko, 2017) was conducted to ensure that the majority of correlations were between 0.2 and 0.8. Second, the Kaiser-Meyer-Olkin (*KMO*) measure of sampling adequacy was calculated using the R package ‘psych’ (version 2.0.9; Kaiser, 1974; Revelle, 2020). *KMO* values above 0.8 were considered *meritorious* while values above 0.9 were considered *marvelous* (Field, 2013). Finally, Bartlett’s test of sphericity was measured to ensure relatedness between items (Bartlett & Fowler, 1937). Bartlett’s test measures whether the item correlation matrix is significantly different from an identity matrix. While a statistically significant result is generally obtained, especially with a large sample size, the lack of significance is an indication of a factorability problem (Field, 2013). Normality was examined using the ‘psych’ package (Revelle, 2020). Univariate normality or non-normality was determined by examining item skew and kurtosis. Item skew and kurtosis below |2.0| were considered normal (Kim, 2013; Knekta et al., 2019). Multivariate normality or non-normality was determined by visual examination of a QQ plot.

The ‘psych’ package was used to compute the EFA (Revelle, 2020). Considering the non-normal nature of the data (see Results) the principal axis factoring method was used for factor extraction in the EFA with an oblique rotation (oblimin; Costello & Osborne, 2005; Knekta et al., 2019; Watson, 2017). Theoretical considerations, examination of the scree plot, and parallel analysis were used to determine the proper number of factors to extract.

The R package ‘lavaan’ (version 0.6-7; Rosseel, 2012) was used to conduct the CFAs with the robust maximum likelihood estimator due to the continuous nature and slight non-normality of the data (Knekta et al., 2019). We used the root-mean-square error of approximation (*RMSEA*; absolute fit index; adequate fit < 0.1), the standardized root-mean-square residual (*SRMR*; absolute fit index; adequate fit <0.08), the comparative fit index (*CFI*; incremental fit index; adequate fit > 0.9), and the Tucker-Lewis index (*TLI*; incremental fit index; adequate fit > 0.9) to evaluate model fit (Browne & Cudeck, 1992; Hu & Bentler, 1999; Knekta et al., 2019; Oh et al., 2013; Tucker & Lewis, 1973). The chi-square value was also calculated. However, chi-squares are quite sensitive to sample size, and a large sample size can result in rejection of an adequate model (Knekta et al., 2019). We calculated the *^2^*/*df* and considered values less than 2 as adequate (Oh et al., 2013). Additionally, we conducted simulations using the ‘dynamic’ package (version 1.1.0; McNeish & Wolf, 2020), which provides dynamic fit index cut-off for different levels, according to the model specified. Marsh and colleagues (2004) caution that strict adherence to goodness-of-fit index cut-offs can result in the rejection of valid models. Therefore, while model fit indices were assessed, they were considered in conjunction with theoretical interpretability when presenting a final model.

To assess the validity evidence for the factor structure for different populations, we performed a series of separate CFA models for six subgroups: men, women, minority, white, high SES, and low SES. Further, we assessed configural, metric, scalar, and conservative measurement invariance (as appropriate; Rocabado et al., 2020) for gender, race, and SES. For these analyses, students who responded “Prefer not to answer” or did not reply to the specific demographic items were excluded from analysis. We defined ‘minority’ as any non-white student, including those who reported belonging to multiple races. School SES was estimated using state reporting of “percent economically disadvantaged.” These values were z-standardized. Schools with z-scores greater than 0 were considered low SES, while those with z-scores less than 0 were considered high SES.

Criterion-related validity was assessed through a series of correlations of latent variables from SASS with external variables. Student latent variable scores were obtained by averaging item responses for each construct. Spearman correlations of the latent variable scores with self-reported STEM GPA were calculated. Students reported individual grades for math, biology, chemistry, and physics, which were converted to standard GPA scores (1 = D, 2 = C, 3 = B, and 4 = A) and averaged to arrive at the STEM GPA score. Additionally, Spearman correlations between school SES and the latent variable scores were assessed. Prior to calculating correlations, distributions were examined for normality using QQ plots, and outliers surpassing 3 standard deviations were excluded. This resulted in the removal of 5 outliers for STEM GPA, and no outliers for school SES.

### Reliability Testing

Internal consistency of each scale was assessed using two methods. First, Cronbach’s alpha was determined using the ‘psych’ package (Revelle, 2020), as this is the statistic most widely accepted for reliability measures. However, several researchers caution against the reliance on Cronbach’s alpha for assessing instrument reliability due to several limitations of the method (Boateng et al., 2018; Knekta et al., 2019; Peters, 2014). Therefore, the omega coefficient, which has been suggested as a replacement for alpha (McNeish, 2018; Peters, 2014), was calculated using the ‘semTools’ package (version 0.5-3; Jorgensen et al., 2020).

Test-retest reliability of SASS was examined using a subset of 62 student responses collected from two schools. Teachers were asked to administer the survey twice in the classroom 7 to 10 days apart, however the time difference in administration ranged from 4 to 12 days. Bland-Altman plots were created using the ‘blandr’ package (version 0.5.1; Datta, 2017) to visually examine the agreement between the first and second measurement of the SASS (Bland & Altman, 1986). Additionally, we examined test-retest reliability by calculating type 3 intraclass correlation coefficients (*ICC(3,1)*; two-way, mixed model, single measures; Shrout & Fleiss, 1979) for each of the 40 items and the six factor scores. The *ICC(3,1)* values and confidence intervals were calculated using the ‘psych’ package. This method is recommended for determining the agreement between two measurements taken using a self-report survey with the same subjects under the same conditions (Koo & Li, 2016; Portney & Watkins, 2015). Following the recommendations of Koo and Li (2016), we considered the following rating scale for *ICC(3,1)* values: *poor* - less than 0.5, *moderate* - 0.5 to 0.75, *good* - 0.75 to 0.90, and

*excellent* - greater than 0.9.

## Results

### Exploratory Factor Analysis

To determine the dimensionality of the new instrument, we performed an EFA using responses from 400 students. Item means and standard deviations for this sample are provided in Table S3. The data exhibited good factorability (*KMO* = 0.9; Bartlett’s *K-squared* = 357.55, *df* = 39, *p* < 10^-15^) and the inter-item correlation matrix exhibited several correlations above 0.3 (Figure S1). All items had skewness and kurtosis below |2.0|, except for two items: “If I choose to pursue a science, technology, engineering, and/or math (STEM) related career, I will be supported in this decision by my family members” (kurtosis = 2.9) and “A scientist is a person who has an unhappy marriage” (kurtosis = 3.2). Examination of the QQ plot suggested possible non-normality of the data. With consideration of the items with non-normal kurtosis and the results from the QQ plot, we proceeded with the principal axis factoring extraction method, which is robust to slightly non-normal data (Knekta et al., 2019).

In addition to our two new latent variables (Negative Perceptions of Scientists and Career Awareness), SASS included items from four constructs from the engineering SCCT (with modifications and additions relevant to high school students and STEM): self-efficacy, outcome expectations, interests, and social supports (Lent et al., 2005). As noted in the Methods, during instrument development, we observed that items regarding expectations of social support were highly correlated with the Outcome Expectations items, suggesting those items could be combined with Outcome Expectations to form a five factor solution (corresponding to Self-Efficacy, Outcome Expectations, Interests, Negative Perceptions of Scientists, and Career Awareness).

However, examination of the scree plot suggested six to seven factors based on the inflection point (Figure S2), and parallel analysis suggested an optimal solution that specified eight factors. Five factors exhibited eigenvalues above 1. Considering these results, and our theoretical expectations (we hypothesized a five factor solution), we performed three EFAs, one with five factors, one with six factors, and one with seven factors. Theoretical considerations based on a priori knowledge, along with examination of the pattern matrices, were used to evaluate and compare model fit for the three models (Table 3, Table S4, Table S5, and Figure S3).

**Table 3.** Item characteristics for six factor EFA and CFA

|  |  | EFA Results (*n* = 400) |  |  |  |  |  |  |  |  | CFA Results (*n* = 532) |  |  |  |  |  |
| --- | --- | --- | --- | --- | --- | --- | --- | --- | --- | --- | --- | --- | --- | --- | --- | --- |
| Category | Item | PA5 | PA4 | PA1 | PA6 | PA3 | PA2 | *h2* | *u2* | *com* | SEE | SEA | OE | IN | SP | CA |
| SEE | Q3_1 | **0.32** | **0.36** | 0.05 | -0.07 | 0.03 | 0.13 | 0.35 | 0.65 | 2.39 | 0.58 |  |  |  |  |  |
| SEE | Q3_2* | **0.44** | **0.53** | 0.08 | -0.14 | 0.03 | 0.08 | 0.65 | 0.35 | 2.21 | 0.66 |  |  |  |  |  |
| SEE | Q3_3 | **0.64** | 0.00 | 0.06 | 0.21 | -0.09 | 0.02 | 0.63 | 0.37 | 1.27 | 0.73 |  |  |  |  |  |
| SEE | Q3_4 | **0.66** | 0.08 | 0.06 | -0.02 | -0.05 | -0.01 | 0.51 | 0.49 | 1.06 | 0.68 |  |  |  |  |  |
| SEE | Q3_5 | **0.64** | 0.01 | 0.01 | -0.05 | -0.08 | 0.03 | 0.43 | 0.57 | 1.05 | 0.69 |  |  |  |  |  |
| SEE | Q3_6 | **0.73** | 0.01 | 0.05 | 0.18 | 0.00 | 0.04 | 0.73 | 0.27 | 1.13 | 0.79 |  |  |  |  |  |
| SEA | Q9_1 | 0.11 | **0.49** | 0.08 | 0.29 | -0.03 | -0.09 | 0.50 | 0.50 | 1.91 |  | 0.71 |  |  |  |  |
| SEA | Q9_2 | 0.00 | **0.64** | -0.01 | 0.24 | -0.07 | -0.03 | 0.53 | 0.47 | 1.29 |  | 0.78 |  |  |  |  |
| SEA | Q9_3 | -0.05 | **0.75** | -0.01 | 0.17 | 0.03 | -0.03 | 0.61 | 0.39 | 1.12 |  | 0.76 |  |  |  |  |
| SEA | Q9_4 | -0.04 | **0.72** | 0.11 | -0.16 | -0.01 | 0.01 | 0.55 | 0.45 | 1.15 |  | 0.60 |  |  |  |  |
| SEA | Q10_1* | **0.31** | **0.31** | 0.09 | 0.22 | -0.06 | 0.11 | 0.54 | 0.46 | 3.37 |  | 0.67 |  |  |  |  |
| SEA | Q10_2* | 0.13 | **0.61** | 0.06 | -0.08 | -0.02 | 0.13 | 0.50 | 0.50 | 1.24 |  | 0.63 |  |  |  |  |
| OE | Q12_1 | 0.09 | 0.10 | **0.61** | -0.05 | 0.06 | 0.04 | 0.46 | 0.54 | 1.14 |  |  | 0.67 |  |  |  |
| OE | Q12_2 | 0.08 | -0.02 | **0.67** | 0.02 | 0.12 | 0.01 | 0.47 | 0.53 | 1.10 |  |  | 0.65 |  |  |  |
| OE | Q12_3 | 0.24 | -0.06 | **0.45** | 0.22 | -0.05 | 0.00 | 0.49 | 0.51 | 2.11 |  |  | 0.66 |  |  |  |
| OE | Q12_4 | 0.17 | -0.08 | **0.53** | 0.13 | -0.02 | -0.03 | 0.43 | 0.57 | 1.41 |  |  | 0.69 |  |  |  |
| OE | Q12_5 | 0.07 | -0.07 | **0.79** | -0.02 | 0.04 | -0.01 | 0.61 | 0.39 | 1.04 |  |  | 0.73 |  |  |  |
| OE | Q12_6 | 0.10 | -0.04 | **0.77** | -0.06 | 0.05 | 0.00 | 0.58 | 0.42 | 1.06 |  |  | 0.71 |  |  |  |
| OE | Q13_1 | -0.09 | 0.08 | **0.52** | 0.16 | -0.09 | 0.10 | 0.45 | 0.55 | 1.46 |  |  | 0.68 |  |  |  |
| OE | Q13_2 | -0.19 | 0.11 | **0.70** | 0.10 | -0.10 | -0.05 | 0.57 | 0.43 | 1.30 |  |  | 0.66 |  |  |  |
| OE | Q13_3 | -0.15 | 0.08 | **0.76** | -0.01 | -0.12 | 0.04 | 0.62 | 0.38 | 1.16 |  |  | 0.70 |  |  |  |
| OE | Q13_4 | 0.13 | 0.10 | **0.51** | 0.10 | -0.05 | 0.07 | 0.50 | 0.50 | 1.36 |  |  | 0.62 |  |  |  |
| IN | Q14_1 | -0.05 | 0.06 | 0.11 | **0.59** | -0.03 | 0.00 | 0.43 | 0.57 | 1.11 |  |  |  | 0.63 |  |  |
| IN | Q14_2 | 0.06 | 0.01 | 0.06 | **0.67** | -0.05 | 0.04 | 0.57 | 0.43 | 1.05 |  |  |  | 0.77 |  |  |
| IN | Q14_3 | 0.09 | -0.02 | 0.10 | **0.65** | -0.02 | 0.17 | 0.67 | 0.33 | 1.23 |  |  |  | 0.81 |  |  |
| IN | Q14_4 | 0.13 | 0.05 | -0.03 | **0.60** | 0.06 | 0.10 | 0.48 | 0.52 | 1.20 |  |  |  | 0.75 |  |  |
| IN | Q14_5 | 0.10 | 0.16 | 0.05 | **0.57** | 0.00 | 0.12 | 0.58 | 0.42 | 1.33 |  |  |  | 0.79 |  |  |
| SP | Q29_1 | 0.05 | -0.06 | -0.01 | 0.00 | **0.86** | -0.02 | 0.75 | 0.25 | 1.02 |  |  |  |  | 0.87 |  |
| SP | Q29_2 | -0.07 | 0.04 | 0.04 | 0.04 | **0.82** | 0.01 | 0.66 | 0.34 | 1.03 |  |  |  |  | 0.78 |  |
| SP | Q29_3 | -0.03 | 0.03 | -0.01 | 0.00 | **0.90** | 0.02 | 0.82 | 0.18 | 1.01 |  |  |  |  | 0.90 |  |
| CA | Q16_1 | 0.12 | -0.08 | 0.18 | 0.11 | -0.04 | **0.40** | 0.35 | 0.65 | 1.89 |  |  |  |  |  | 0.42 |
| CA | Q16_2 | 0.05 | -0.16 | -0.05 | 0.01 | -0.04 | **0.54** | 0.30 | 0.70 | 1.21 |  |  |  |  |  | 0.47 |
| CA | Q16_3 | 0.01 | -0.11 | -0.01 | 0.02 | 0.03 | **0.66** | 0.43 | 0.57 | 1.06 |  |  |  |  |  | 0.55 |
| CA | Q16_4 | 0.03 | 0.05 | -0.01 | -0.08 | 0.05 | **0.55** | 0.30 | 0.70 | 1.09 |  |  |  |  |  | 0.54 |
| CA | Q16_5 | -0.01 | 0.11 | 0.02 | -0.12 | 0.03 | **0.66** | 0.42 | 0.58 | 1.13 |  |  |  |  |  | 0.59 |
| CA | Q16_6 | 0.03 | 0.01 | 0.01 | 0.08 | -0.05 | **0.72** | 0.61 | 0.39 | 1.04 |  |  |  |  |  | 0.82 |
| CA | Q16_7 | -0.08 | 0.09 | -0.01 | 0.00 | -0.02 | **0.83** | 0.68 | 0.32 | 1.04 |  |  |  |  |  | 0.80 |
| CA | Q16_8 | -0.07 | -0.03 | -0.02 | 0.08 | -0.02 | **0.66** | 0.44 | 0.56 | 1.06 |  |  |  |  |  | 0.68 |
| CA | Q16_9 | 0.06 | -0.05 | 0.11 | 0.11 | 0.04 | **0.58** | 0.46 | 0.54 | 1.19 |  |  |  |  |  | 0.62 |
| CA | Q16_10 | 0.05 | -0.09 | -0.04 | 0.04 | 0.03 | **0.66** | 0.45 | 0.55 | 1.07 |  |  |  |  |  | 0.63 |

The five factor solution explained 50% of the variance in the model. Each of the factors had a sum of squared loadings above 2, indicating that they were contributing positively to the model. All cross factor loadings were below 0.30, except for three items. These were equal to or below 0.40 (“I am confident in my ability to succeed in a career that requires knowing science and using math”, “I feel prepared to take college courses that use science knowledge”, and “Having a career that uses science, technology, engineering, and/or math (STEM) will allow me to do exciting work”). All three of these items cross-loaded on the a priori expected Interests factor. Furthermore, four items which we expected to load on the Self-Efficacy factor instead loaded on Interests above 0.50 (“I am confident in my ability to succeed in a career that requires: - informing people about topics in science and/or medicine, -figuring out how things work, -doing lab work or experiments, and -talking about science”). Pattern coefficients were above 0.50 for 30 of the 40 items. All items in the Interests factor loaded poorly, with pattern coefficients below 0.45. One item (“Rate your interest in: thinking about topics that relate to my health”) did not load above 0.27 on any factor. In sum, these results suggested that the addition of a sixth factor to account for the four Self-Efficacy items loading with Interests may improve the model.

The six factor model explained a greater percentage of the variance (53%) than the five factor model and all factors had a sum of squared loadings above 2. The four Self-Efficacy items that unexpectedly loaded with the Interests factor in the five factor model comprised a new factor when a six factor model was examined. Based on examination of the item wording, we chose to call this new factor Self-Efficacy-Experience and categorized the remaining Self-Efficacy items as Self-Efficacy-Academic. Three items were cross-loaded in this six factor model (“I am confident in my ability to succeed in a career that requires: -understanding or interpreting graphs, - knowing science and using math,” and “I feel prepared to take college courses that use science knowledge”) and they had roughly equivalent pattern coefficients for the a priori Self-Efficacy-Academic factor and Self-Efficacy-Experience. We chose to classify the first two items as Self-Efficacy-Experience and the third as Self-Efficacy-Academic based on our evaluation of the item wording. All items had pattern coefficients above 0.50, except for five, which were above 0.30. Of the 40 items in the instrument, 25 exhibited pattern coefficients above 0.60.

The seven factor model explained a greater percentage of the variance (55%) than the five or six factor models. However, only six of the seven factors had a sum of squared loadings above 2. The new factor, which corresponded with three of four social support items, had a sum of squared loadings of 1.91. The three items that were cross-loaded in the six factor model were again cross-loaded in the seven factor model. An additional three items were cross-loaded at 0.3 or above (“If I choose to pursue a science, technology, engineering, and/or math (STEM) related career, I will: -be supported in this decision by my family members, -feel that my friends and/or family are proud,” and “Rate your interest in thinking about topics that relate to my health”). All items had pattern coefficients above 0.50, except for seven, which were above 0.30. Of the 40 items in the instrument, 23 exhibited pattern coefficients above 0.60. Considering the increased number of cross-loaded items, we did not continue examination of a seven factor structure in the confirmatory factor analyses.

### Confirmatory Factor Analysis

Following EFA, CFA was performed using an independent sample of 532 student responses. Item means and standard deviations for this sample are provided in Table S3. The data exhibited good factorability (*KMO* = 0.9; Bartlett’s *K-squared* = 602.23, *df* = 39, *p* < 10^-15^) and the inter-item correlation matrix exhibited several correlations above 0.30 (Figure S4). All items had skewness and kurtosis below |2.0|, except for three items: “If I choose to pursue a science, technology, engineering, and/or math (STEM) related career, I will be supported in this decision by my family members” (kurtosis = 2.2), “A scientist is a person who has an unhappy marriage” (kurtosis = 2.5), and “A scientist is a person who is lonely” (kurtosis = 2.0). As seen with the EFA sample, examination of the QQ plot suggested slight non-normality of the data. With consideration of the items with non-normal kurtosis and the results from the QQ plot, we proceeded with the robust maximum likelihood estimation method due to the continuous nature and the slight non-normality of the data (Knekta et al., 2019). For all models tested, all item estimates were significant (*p* < 10^-3^).

We began by constructing five factor and six factor models of the data (Figure 1, Table 3, and Table S6). A single factor model was also calculated and is provided for reference (Table S7). Fit indices for the five factor and six factor model were similar, with *CFI* and *TLI* values between 0.7 and 0.8, *RMSEA* of 0.08, and *SRMR* of 0.07 (Table 4). Neither of the models exhibited a *χ^2^*/*df* less than 2 (the suggested cut-off), but this may be because the measure is inaccurate for larger sample sizes (Knekta et al., 2019). While the *CFI* and *TLI* values were below the cut-off for an adequate fit (0.9), the *RMSEA* was below 0.1, indicating an adequate fit by that measure (Browne & Cudeck, 1992; Hu & Bentler, 1999; Knekta et al., 2019; Oh et al., 2013; Tucker & Lewis, 1973). Additionally, the *SRMR* was below 0.08, indicating acceptable model fit (Hu & Bentler, 1999). Dynamic fit index cut-offs are provided in Table S8. For both models, all item loadings were between 0.50 and 0.90, with two exceptions: Career Awareness items regarding doctor (five factor: 0.41; six factor: 0.42) and animal care technician (five factor: 0.47; six factor: 0.47). Factor loadings were minimally changed for most items when comparing the five and six factor models, with the exception of all items in the Self-Efficacy-Experience, and Self-Efficacy-Academic factors, which generally exhibited higher loading scores in the six factor model (average difference in item loading = 0.06). In sum, the CFA and EFA results both suggest that the six factor model is a better fit for the data.

**Figure 1.**
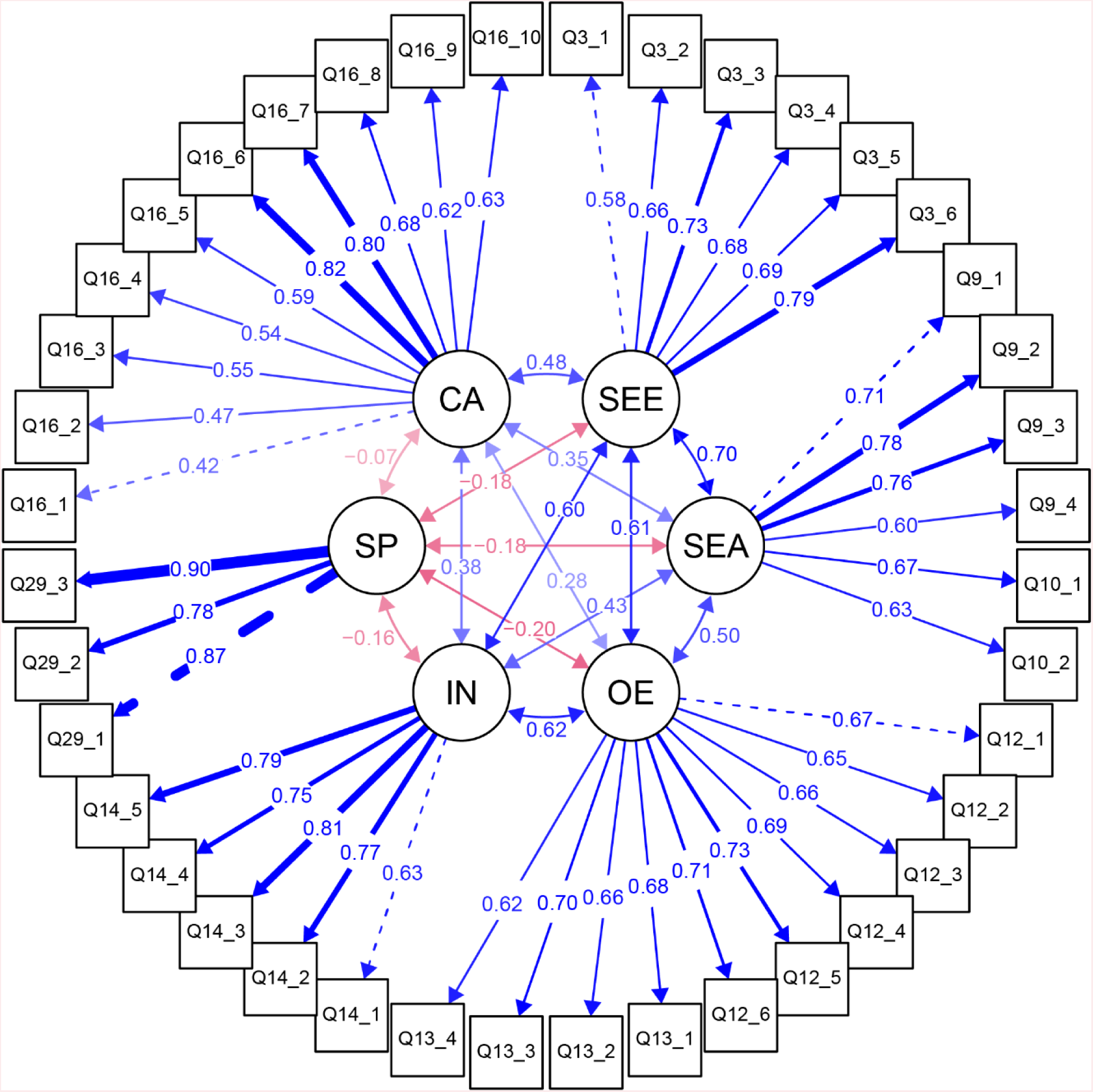
Confirmatory factor analysis item loadings and factor correlations Results from the six factor CFA model are shown. Factors are represented by circles, while survey items are represented by squares (see Table 1 for item descriptions). The one-directional arrows from the factors to the items represent the standardized factor loadings; the arrows are weighted by the strength of the factor loading and the dashed arrows indicate which item estimates were fixed to 1 in the model. The bi-directional arrows between factors represent the correlations between the factors. Blue arrows indicate a positive loading or correlation. Red arrows indicate a negative loading or correlation. All item estimates were significant at *p* < 10^-3^. Latent variable correlations were significant at *p* < 10^-2^, except for Negative Perceptions of Scientists with Career Awareness, which was *p* = 0.20. Abbreviations: SEE = Self-Efficacy-Experience, SEA = Self-Efficacy-Academic, OE = Outcome Expectations, IN = Interests, SP = Negative Perceptions of Scientists, CA = Career Awareness.

**Table 4.** CFA fit indices

| Model | *n* | $\chi^2$ | *df* | $\chi^2/df$ | *CFI* | *TLI* | *RMSEA* | *SRMR* |
| --- | --- | --- | --- | --- | --- | --- | --- | --- |
| One factor | 532 | 6999 | 740 | 9.5 | 0.45 | 0.42 | 0.13 | 0.12 |
| Five factor | 532 | 3506 | 730 | 4.8 | 0.76 | 0.75 | 0.08 | 0.07 |
| Six factor | 532 | 3231 | 725 | 4.5 | 0.79 | 0.77 | 0.08 | 0.07 |
| Reduced six factor | 532 | 2057 | 614 | 3.35 | 0.83 | 0.81 | 0.07 | 0.06 |
| Men | 248 | 1382 | 614 | 2.3 | 0.80 | 0.79 | 0.08 | 0.07 |
| Women | 269 | 1323 | 614 | 2.2 | 0.84 | 0.82 | 0.07 | 0.07 |
| White | 257 | 1356 | 614 | 2.2 | 0.83 | 0.82 | 0.07 | 0.07 |
| Minority | 191 | 1405 | 614 | 2.3 | 0.76 | 0.74 | 0.09 | 0.08 |
| High SES | 184 | 1372 | 614 | 2.2 | 0.77 | 0.75 | 0.09 | 0.08 |
| Low SES | 265 | 1392 | 614 | 2.3 | 0.84 | 0.82 | 0.07 | 0.07 |

For *CFI*, *TLI*, *RMSEA*, and *SRMR*, robust measures were used. Abbreviations: *n* = number of students, $\chi^2$ = chi-square, *df* = degrees of freedom, *CFI* = comparative fit index, *TLI* = Tucker-Lewis index, *RMSEA* = root-mean-square error of approximation, and *SRMR* = standardized root-mean-square residual.

Correlations between latent factors for the six factor model are presented in Table 5. The latent factor correlations range from -0.2 (Negative Perceptions of Scientists with Self-Efficacy-Academic, Self-Efficacy-Experience, Outcome Expectations, and Interests) to 0.7 (Self-Efficacy-Experience with Self-Efficacy-Academic). In general, the SCCT latent variables (Self-Efficacy-Experience, Self-Efficacy-Academic, Outcome Expectations, and Interests) were highly correlated with each other. The Career Awareness factor exhibited moderate correlation with the SCCT factors, and Negative Perceptions of Scientists was weakly correlated with these factors.

**Table 5.** CFA six factor model latent variable correlations

|  | SEE | SEA | OE | IN | SP |
| --- | --- | --- | --- | --- | --- |
| SEA | 0.70* | - |  |  |  |
| OE | 0.61* | 0.50* | - |  |  |
| IN | 0.60* | 0.43* | 0.62* | - |  |
| SP | -0.18* | -0.18* | -0.20* | -0.16* | - |
| CA | 0.48* | 0.35* | 0.28* | 0.38* | -0.07 |

\* $p < 10^{-2}$ . Abbreviations: SEE = Self-Efficacy-Experience, SEA = Self-Efficacy-Academic, OE = Outcome Expectations, IN = Interests, SP = Negative Perceptions of Scientists, CA = Career Awareness.

### Refining the Model

With the goal of constructing a better performing CFA model, we revisited the six factor EFA results (Table 3). Items were identified for elimination based on cross-loading and theoretical considerations. We deleted items in a stepwise fashion, resulting in a final model with three items removed: I am confident in my ability to succeed in a career that requires knowing science and using math (Q3_2), I feel prepared to take college courses that use science knowledge (Q10_1), and I feel prepared to take college courses that use math skills (Q10_2). Items Q3_2 and Q10_2 were both cross-loaded on Self-Efficacy-Experience and Self-Efficacy-Academic and therefore were primary candidates for dropping from the model. Item Q10_1 was additionally considered for removal based on its content pairing with Q10_2. Further, items Q10_1 and Q10_2 both require students to consider a *future* academic trajectory (i.e. going to college), whereas the traditional self-efficacy items are focused on a *present* academic path (i.e. attaining high grades in high school classes). Therefore, we suggest these two items may reflect a different theoretical meaning than that for the self-efficacy items in the engineering survey. Finally, item Q3_1 (I am confident in my ability to succeed in a career that requires understanding or interpreting graphs) was also cross-loaded. However, we left this item in the model because it is a crucial STEM skill and a fundamental part of the latent variable. Further, this item is useful when measuring the impact of educational interventions that specifically target graph reading skills.

Results for the reduced six factor EFA and CFA are reported in Table 4, Table S9, and Table S10. As expected, the reduced six factor model produced good EFA results, with the exception of one item with cross-loading (Q3_1). All item loadings were above 0.4, except for “Doctor” (Q16_1), which loaded on the a priori expected Career Awareness factor at 0.39. Of the 37 items in the reduced factor model, 25 exhibited pattern coefficients above 0.60. Overall, this reduced six factor model performed similarly in EFA to a model in which all items were retained.

CFA fit indices for the reduced six factor model were slightly improved when compared with the model retaining all items (Table 4). Measures of incremental fit, *CFI* and *TLI*, improved to 0.83 and 0.81, respectively. The absolute fit metrics of *RMSEA* (0.07) and *SRMR* (0.06) support the notion of structural validity for this model. The RMSEA and SRMR exceeded level 5 dynamic fit index cut-offs (RMSEA < 0.05, SRMR < 0.07), but the CFI did not (CFI > 0.89). All item loadings were between 0.50 and 0.90, with the same two exceptions as exhibited in the five and six factor models reported above. In sum, the CFA and EFA results both suggest that the reduced 37-item six factor model is a slightly better fit for the data than the 40-item six factor model in which all items were retained. We performed all subsequent analyses with the reduced six factor model.

### Measurement Invariance

Confirmation of the structural validity in subgroups is often used to support construct validity (e.g., Unfried et al., 2015). Therefore, we performed CFAs with subsamples of the data, split by gender, race, and SES. Models for all subsamples performed similarly (Table 4; *CFI* and *TLI* between 0.7 and 0.8; *RMSEA* between 0.07 and 0.09; *SRMR* between 0.07 and 0.08). Consistent with their smaller sample sizes, these subsample models exhibited *χ^2^*/*df* closer to 2 than seen with the full sample set. We followed examination of these baseline models with measurement invariance tests for each demographic variable (Table S11). Metric invariance was achieved for gender and SES, while scalar invariance was achieved for race. These results indicate that the six factor model structure is acceptable and stable across gender, race, and SES in our sample. Additionally, the invariance testing results suggest SASS factor means can be compared between minority and white students.

### Criterion Validity

One method of gauging the meaningfulness of an instrument’s factors is to test the relationship of each latent variable within the instrument to an independent criterion. We compared self-reported STEM GPA to Self-Efficacy-Experience scores and Self-Efficacy-Academic scores, which measure students’ confidence in using STEM-related skills or their confidence in their ability to obtain a B or better in STEM classes, respectively (Figure 2A-B). Considering the shared relationship of STEM GPA, Self-Efficacy-Experience, and Self-Efficacy-Academic to academic performance, we hypothesized that both Self-Efficacy-Experience and Self-Efficacy-Academic scores would increase with students’ self-reported STEM GPAs. As expected, STEM GPA was positively correlated with Self-Efficacy-Experience and Self-Efficacy-Academic scores in our sample (Self-Efficacy-Experience: *rho* = 0.30, *p* < 10^-11^; Self-Efficacy-Academic: *rho* = 0.62, *p* < 10^-15^). A meta-analysis of 121 independent samples found a correlation between academic achievement and academic interest (Schiefele et al., 1992).

**Figure 2.**
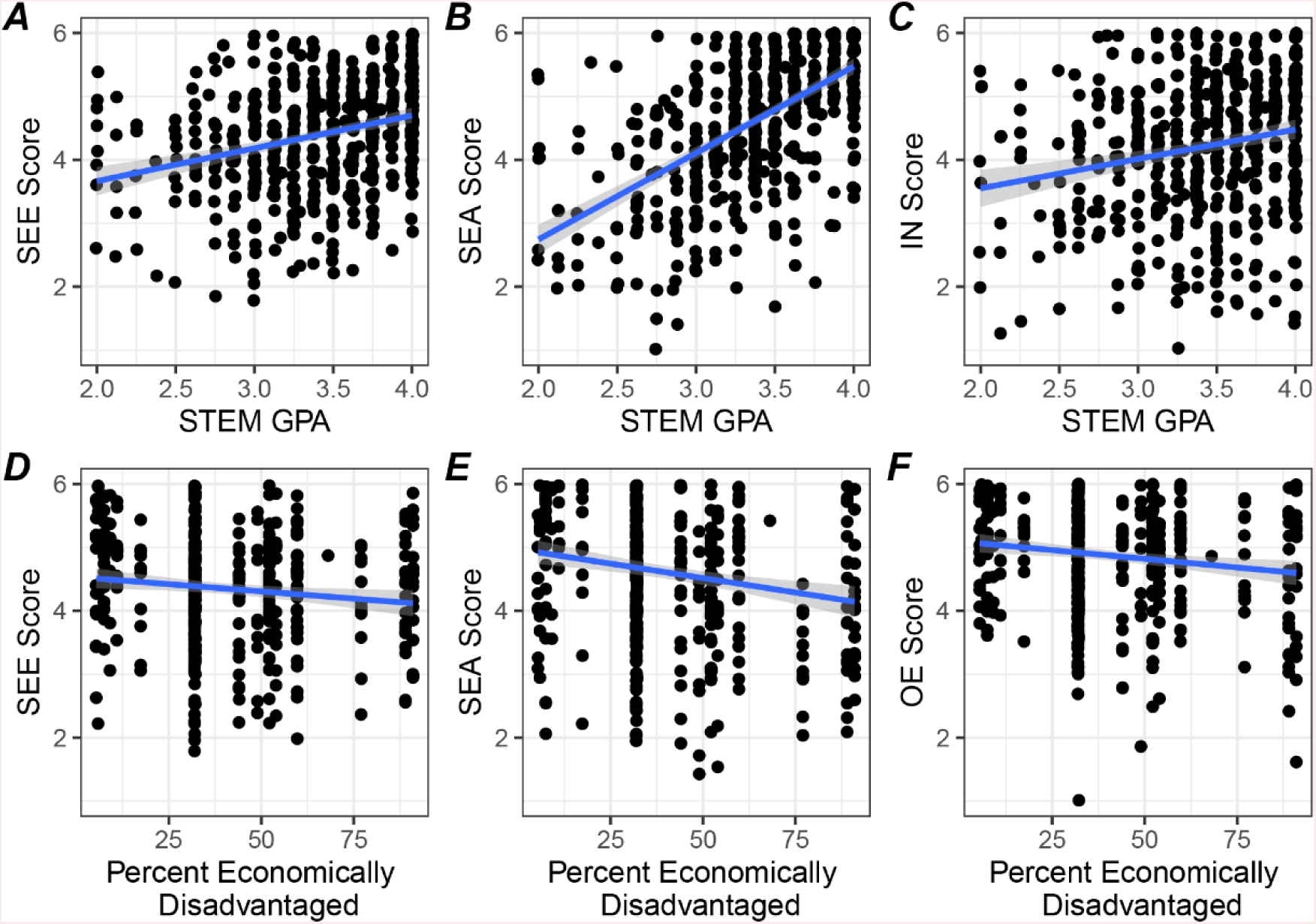
Criterion-related validity testing Scatter plots illustrating the relationship of factor scores and external variables are shown with linear regression lines. Spearman’s correlations and *p* values were also calculated for SASS factors with STEM GPA: A) Self-Efficacy-Experience score (*rho* = 0.30, *p* < 10^-11^); B) Self-Efficacy-Academic score (*rho* = 0.62, *p* < 10^-15^); C) Interests score (*rho* = 0.21, *p* < 10^-5^); and for SASS factors with school percent economically disadvantaged: D) Self-Efficacy-Experience score (*rho* = -0.11, *p* = 10^-2^); E) Self-Efficacy-Academic score (*rho* = -0.17, *p* < 10^-3^); and F) Outcome Expectations score (*rho* = -0.14, *p* < 10^-2^). A high percentage of economically disadvantaged students corresponds with low SES for the school. Because the SES data is collected at the school level, the values are clustered at discrete values on the horizontal axis. Abbreviations: SEE = Self-Efficacy-Experience, SEA = Self-Efficacy-Academic, OE = Outcome Expectations, IN = Interests, SP = Negative Perceptions of Scientists, CA = Career Awareness.

Therefore, we examined the relationship between student STEM GPA and Interests scores (Figure 2C). As expected, a statistically significant correlation was found (*rho* = 0.21, *p* < 10^-5^). Previous research suggested a relationship between SES and self-efficacy (Navarro et al., 2007), as well as SES and outcome expectations (Turner et al., 2019). Therefore, we examined the correlation between SES and Self-Efficacy- Experience, Self-Efficacy-Academic, and Outcome Expectations scores (Figure 2D-F). Our results indicated that a lower percent economically disadvantaged (higher SES) was associated with higher Self-Efficacy-Experience (*rho* = -0.11, *p* = 10^-2^), Self- Efficacy-Academic (*rho* = -0.17, *p* < 10^-3^), and Outcome Expectations (*rho* = -0.14, *p* < 10^-2^) scores. For completeness, we examined the correlation of STEM GPA and SES with the remaining latent variables (Figure S5). No correlations were significant using a *p* < 10^-2^ threshold, except STEM GPA with Outcome Expectations score (*rho* = 0.24, *p* < 10^-7^). Taken together, these results suggest that these factors can be interpreted with good criterion validity.

### Reliability

Internal consistency of an instrument is a measure of how well each of the items in a latent variable relate to each other. We evaluated the internal consistency of the instrument by measuring Cronbach’s alpha, the “gold standard,” and omega for each of the six constructs. Ideally the alpha and omega values should be greater than 0.70. However, values that are greater than 0.95 would be indicative of high redundancy in the instrument items (Tavakol & Dennick, 2011). For SASS (using the reduced item variant), alpha and omega values were 0.8 for Self-Efficacy-Experience and Self-Efficacy-Academic, and 0.9 for Outcome Expectations, Interests, Negative Perceptions of Scientists, and Career Awareness. Therefore, the instrument demonstrated *good* to *excellent* internal consistency.

While many SCCT instruments are used to take a snapshot of participant views, we planned to use SASS for evaluating the impact of a five day classroom intervention on student attitudes related to STEM careers. Therefore, it was important to assess the stability of the instrument over time. Repeated measures invariance testing can evaluate how equivalently the items in the measure function across time points, but the subset of our sample for which we had repeated scores was too small (*n* = 62) to support these analyses. With a larger sample, measurement invariance analyses could provide evidence for how stable the model is over time. Instead, we used *ICC(3, 1)* to assess the test-retest reliability of SASS as a first step in determining reliability of the instrument in our population over time (Table S12). Students in the sample subset took the survey an average of nine days apart. The *ICC(3,1)* values were *good* for Self-Efficacy-Academic (0.90), *moderate* for Self-Efficacy-Experience, Outcome Expectations, Interests, and Negative Perceptions of Scientists (0.64, 0.53, 0.57, and 0.72, respectively), and *poor* for Career Awareness (0.47). Bland-Altman plots are presented in Figure S6.

## Discussion

### Validation and Reliability of SASS

With the goal of understanding what factors influence career choice in high school students, we developed the SASS questionnaire, which measures attitudes surrounding STEM careers. When used with a high school cohort, the instrument can be used reliably, and there is evidence for good structural and criterion validity. In developing SASS, hard cut-offs for the goodness-of-fit indices were carefully considered in conjunction with theory and content validity when determining model fit. Although the goodness-of-fit measures obtained for the CFA models (*^2^*/*df*, *CFI*, *TLI*, and *RMSEA*) did not universally exceed standard cut-offs, we note that these cut-offs are meant to be taken as a ‘rule of thumb,’ and not applied without theoretical consideration (Marsh et al., 2004). Our sample included students from a diverse array of schools, and it is possible this high heterogeneity did not allow for superlative goodness-of-fit measures. Previous SCCT survey instruments, which exhibited goodness-of-fit metrics that exceeded the recommended cut-offs, have been evaluated using a more homogenous sample (e.g., Mau et al., 2019; Rasheed Ali & Saunders, 2009; Turner et al., 2019), or using data from students of a single demographic background (e.g., Garriott et al., 2013). Because we wanted to evaluate interventions in a diverse array of classrooms, an instrument that had been evaluated with a heterogeneous sample would be beneficial, even though heterogeneous samples might produce lower fit metrics. Regardless, the *RMSEA* values for all models examined (except the single factor model) were below 0.1, indicating an acceptable fit according to this metric (Browne & Cudeck, 1992). After examination of results from EFA and CFA, as well as consideration of item wording and our study goals, we recommend the use of a six factor model for SASS.

In an effort to produce a model with a better fit to the data, we reviewed results from the six factor EFA and removed three under-performing items in a stepwise fashion. The EFA results from this reduced six factor model were very similar to those obtained with the full sample. The CFA results from this reduced six factor model, slightly improved fit indices for the model. We performed all follow-up analyses using this 37-item instrument and presented this as the final version of SASS.

Criterion validity estimates how well a psychometric construct agrees with an external measure or a theoretical relationship. Using the reduced six factor model, we assessed criterion validity for factors derived from the SASS instrument by examining relationships of the latent variables to self-reported STEM GPA and school SES. Consistent with previous research, we found significant positive relationships between STEM GPA and Self-Efficacy-Experience, Self-Efficacy-Academic, and Interests scores (Lopez & Lent, 1992; Navarro et al., 2007; Schiefele et al., 1992). In examining school SES as a predictor of Self-Efficacy-Experience, Self-Efficacy-Academic, and Outcome Expectations scores, we identified significant relationships between SES and the three SCCT variables, consistent with previous research (Navarro et al., 2007; Turner et al., 2019). While the relationships between SES and these SCCT constructs were significant, they were small. This is consistent with research conducted in Australia that reports relatively small differences in career aspirations related to socio-economic status (Gore et al., 2015). Additionally, we obtained school-level SES data, which obscures differences between students within a school; student-level SES data might provide more accurate estimates of the relationships. Taken together, the correlations of SASS variables with STEM GPA and SES demonstrate the criterion validity of the meaning ascribed to the scores obtained from the instrument.

We found *good* to *excellent* internal consistency of the six latent variables using alpha and omega analyses. This was expected, especially for the Self-Efficacy-Academic, Outcome Expectations, and Interests factors that were based on reliable measures from another instrument (Lent et al., 2005). Test-retest reliability of SASS factors were *good* or *moderate* for five of the six latent variables. The factor with the best test-retest reliability was Self-Efficacy-Academic, suggesting that students’ belief in their ability to perform well academically in the STEM fields was consistent between the two timepoints. The Career Awareness factor displayed the least consistency between the two measurements, exhibiting *poor* test-retest reliability. One explanation for low reliability of this factor may be the assumed dependence on student confidence in how much they know, which could be impacted by their mood or emotional state on any particular day. Additionally, the test-retest reliability was assessed in a small subset of students from two college-preparatory schools and may not reflect the performance of the survey in a broader sample of schools. The *ICC* values are impacted by the variability of a sample; the more homogenous a sample, the lower the *ICC* values (Aldridge et al., 2017). Therefore, it isn’t unreasonable to expect higher *ICC* values in a larger, more heterogeneous sample, and further test-retest reliability testing could be helpful. Finally, the Career Awareness factor may be sensitive to intervening events that improve knowledge about a career or reduce confidence in that knowledge. The Career Awareness factor was designed to measure the effectiveness of interventions, so sensitivity to changing career awareness is crucial.

### Identification of SCCT Variables

The 37-item reduced six factor model robustly identifies three constructs adapted from a similar SCCT instrument for engineering majors (Lent et al., 2005): Self-Efficacy-Academic, Outcome Expectations, and Interests. With the goal of broadening the SASS Self-Efficacy construct, we developed items to measure self-efficacy as it relates to STEM skills. A five factor model, which combined the traditional academic and newly developed skill items into a single factor, was not as coherent in the EFA results. This suggests that perceived STEM skills and perceived academic skills reflect different underlying latent constructs. For example, a student might not feel confident in trigonometry, but might feel capable in figuring out how their bicycle works. We named the new construct *Self-Efficacy-Experience* because the skills items reflect STEM experience outside the classroom. As expected, correlations among these four latent variables (Self-Efficacy-Experience, Self-Efficacy-Academic, Outcome Expectations, and Interests) were high (0.36 to 0.62), confirming the well-established relationship of the SCCT constructs (e.g., Cameron et al., 2015; Lent et al., 2005, 2018; Turner et al., 2019).

A notable SCCT latent variable missing from SASS is a measure of student goals. During instrument development, we tested versions of SASS that included items which comprised a goal setting factor. These items exhibited consistently low coefficients (<0.4), cross-loaded into multiple factors, and did not have high correlations (>0.5) with each other in EFA. After four rounds of rewording questions and piloting new versions with over 1,000 students, we decided to remove the goals measure. Unlike college students, who are thinking about a major and a potential career path, high school students may not have a clear grasp on their goals. Young adolescents may have too many interests to pin down a single goal, they may conflate interests and goals, or they may not be considering their future at all. In a meta-analysis of 41 SCCT studies, 16 out of 16 high school or middle school oriented studies did not include a goal setting measure (Sheu & Bordon, 2017). Therefore, exclusion of a goal setting latent variable is well supported for an instrument aimed at adolescents.

### Inclusion of New Variables

During educational interventions, students may change their perceptions of STEM or of the people working in STEM. This is especially important for high school students who are exploring potential career aspirations, particularly if they are envisioning themselves as scientists. With this in mind, we developed the Negative Perceptions of Scientists latent variable, to measure student perceptions of scientists. While student held perceptions of scientists were previously linked with self-efficacy, interests, and goals (Christidou, 2011; Wyer, 2003), we found small correlations (below |0.2|) between Negative Perceptions of Scientists and the SCCT constructs. Further work, such as path analysis, could elucidate the relationship between student Negative Perceptions of Scientists and other factors in the SCCT framework.

The three items included in Negative Perceptions of Scientists constituted a robust and independent factor, and they conveyed the negative perceptions that scientists are isolated, work-obsessed, and have poor marriages. These stereotypes were based on work from the 1950’s (Mead & Métraux, 1957). Although the items are similar to those found in two independent instruments that measure stereotypes of scientists (Krajkovich & Smith, 1982; Wyer et al., 2010), it is likely that perceptions of scientists have changed in the nearly 70 years that have passed since the initial observations of Mead and Métraux. For example, the stereotype that science is for men has already started to shift, as shown by Scholes and Stahl (2020), who reported that only two out of 45 students recorded scientists as a man, while the majority of students used ungendered language. The addition of items to reflect this change in gender stereotypes could prove valuable.

Preliminary examination of the Negative Perceptions of Scientists item responses indicated that students were extremely consistent in their scoring of the three items. Furthermore, most students scored a 1 or 2 for all three items, indicative of a floor effect. Although these items constitute a mathematically reliable measure with good validity evidence, they may not represent modern views of scientists. In other words, the participants may rate the three items similarly because they are ridiculous statements to contemporary U.S. high school students. Rewording of the current items or addition of new items may improve the factor. Additional items could include examination of scientist stereotypes related to race, class, or gender. Similarly, while the item pertaining to an “unhappy marriage” correlated highly with the other two Negative Perceptions of Scientists items in our sample, the language is not inclusive of all forms of romantic relationships and could be updated to “romantic partnership” or “family life.” In sum, an expanded Negative Perceptions of Scientists measure might shed more light on student perceptions and the impact of classroom interventions aimed at increasing STEM career choice.

The decision to pursue a career in STEM requires more than just traditional SCCT factors, like social support, confidence, and interests. A student must know the career exists and have some idea of what it means to perform the job. We built the Career Awareness items to reflect student knowledge of selected STEM careers. While this factor allows us to capture self-perceived knowledge of careers, it is not an objective measure of student career knowledge, and therefore it might be influenced by their confidence (i.e., self-efficacy). For example, a very insecure student might provide a low rating in knowledge of animal care technician knowledge, even if they had a family member with that job. However, the correlations among Career Awareness and the SCCT variables were low (0.28 to 0.48), suggesting that the influence of confidence on this factor may be small. Nevertheless, self-efficacy is a complex and powerful psychological characteristic (Bandura, 2012), and further research into how SCCT constructs relate to Career Awareness could be informative.

The careers we selected for the Career Awareness factor may not be representative of all STEM careers, but these items could potentially be tailored to an intervention by a given researcher. In support of this view, early versions of SASS included other combinations of careers and we did not observe cross-loading with other factors in the preliminary EFA results. Indeed, the CFA results indicate low correlations with other factors, suggesting that career customization may be possible. Inclusion of a career awareness variable in the SCCT framework will help researchers learn how self-reported career knowledge is impacted when performing educational interventions.

### Limitations

Development of psychometric instruments is difficult, and the evidence for validity is rarely presented without limitations. Generalizability is always a concern; factor analysis results may be different in an independent sample. While the data for this study was obtained exclusively from schools in the northeastern United States, a wide representation of school characteristics (public, private, high SES, and low SES) was included. Additionally, students ranged from 9th to 12th grade, which may represent students with a wide range of attitudes and maturity. Measurement invariance tests in subgroups of our sample population provide validity evidence for the use of the instrument across various demographics. However, the small sample sizes precluded analysis of measurement invariance across grades, so further work could explore differences and similarities between these subgroups, especially if they are to be considered a single population. In addition, the metrics reported in this study may not generalize to dissimilar populations, such as those from other countries or other regions of the United States. Likewise, the test-retest sample size was not large enough to obtain strong measurement invariance statistics, so we do not provide evidence regarding how equivalently the items and factors in the measure are functioning across time points. Instead, our results only demonstrated how similar scores were before and after a delay.

Most of the items included in the instrument were adapted from an established instrument and were assessed for content validity by experts in the field. However, due to limited resources and time, we did not assess how adolescents interpreted the instrument items. Detailed interviews with the target population might provide additional qualitative insight into the consistency of student interpretation. This will likely prove especially useful in further development of the Negative Perceptions of Scientists construct, especially with respect to desirable stereotypes, which were not assessed in this instrument. Further, during instrument development, analyses had suggested the positive perceptions items might form a separate factor; not including positive perceptions could result in construct underrepresentation.

We removed three items from the interim 40-item SASS instrument after reviewing the EFA and CFA results, to produce the final 37-item version presented in this manuscript. Modifying the scale without obtaining a new sample could result in overfitting the model to this particular sample. Further, the scores used to represent latent constructs were obtained using mean values for the component items. While this approach is common, it assumes that the distance between each of the response options is equal. Item response theory analyses could explore this further.

Finally, several limitations surrounding the data collection should be noted. First, the instrument items were presented sequentially (not randomized or shuffled). This was done in order to align with the methods of Lent and colleagues (2005), as SASS was adapted from their instrument. However, order or priming effects could arise; that is, an answer could be influenced by the question that preceded it. Additionally, the authenticity of data collected from self-reported surveys is hard to assess because people may be dishonest. This is especially important to consider with respect to self-reported GPA, which was used to evaluate validity of the constructs. In an effort to collect high quality data, we asked teachers to administer surveys during class time and requested that the surveys were not tied to grades or attendance requirements. Future versions of the instrument could include embedded questions to act as quality control measures of the data similar to Semsar and colleagues (2011).

### Conclusions

In summary, we have created a useful and concise tool for assessing educational interventions intended to impact high school student’s feelings, views, and knowledge about STEM careers. Accurate assessment of these interventions is essential in preventing attrition through the “leaky pipeline.” The SASS instrument extends and modifies SCCT for a high school population with the addition of several factors while still maintaining a short length. We expanded the Self-Efficacy construct to include STEM skills outside of the classroom (Self-Efficacy-Experience), and added two new factors, Career Awareness and Negative Perceptions of Scientists. Career knowledge and negative stereotypes are crucial characteristics in determining STEM career trajectories. To our knowledge, this is the first instrument that has integrated both Career Awareness and Negative Perceptions of Scientists into the SCCT framework. With SASS, researchers can investigate how the SCCT factors interact with career awareness and negative perceptions of scientists, how the factors influence each other, how they predict intention to pursue a STEM career, and how they are influenced by educational interventions.

## Author Contributions

E.K.M., J.H., B.J., and K.S.S. were responsible for the study concept, design, and survey item revision. E.K.M. and J.H. collected the data. E.K.M. and K.S.S. analyzed the data. The paper was written by E.K.M. and K.S.S., with additional input from the other authors. All authors critically reviewed and approved the manuscript for publication.

## Acknowledgments

The authors thank Leslie Schneider, Russel Faux, Revati Masilamani, and David Reider for consultation during the collection of pilot data and the instrument revision process; as well as providing expert review and modification for the question wording. We thank Robert W. Lent for kindly providing a copy of the SCCT survey for engineering majors. The authors thank Nicky Case for providing creative input when naming the survey instrument. Also, reviewers suggested substantial changes, including prompting us to create the reduced 37-item version. Finally, we would like to acknowledge the roles of the teachers and students for making this study possible.

Research reported in this publication was supported by a National Science Foundation Innovative Technology Experiences for Students and Teachers Grant (#1614167) and a National Institute of Health Science Education Partnership Award (#R25GM137369). The content is solely the responsibility of the authors and does not necessarily represent the official views of the National Institutes of Health, the National Science Foundation, the U.S. Department of Veterans Affairs, or the United States Government.

## Supplemental Tables

**Table S1.** Exemplar evolution of item wording

| **SCCT Construct** | **Engineering SCCT<sup>a</sup>** | **Pilot SASS** | **Final SASS** |
| --- | --- | --- | --- |
| Self-efficacy | How confident are you that you could: Continue on in engineering even if you felt that, socially, the environment in these classes was not very welcoming to you. | I am confident in my ability to take science classes that interest me even if I don't "fit in" with other students. | Question dropped due to poor clustering in early exploratory factor analysis. |
| Outcome Expectations | Graduating with a BS degree in engineering will likely allow me to go into a field with high employment demand. | Having a STEM career will allow me to get a job that is in high demand. | Having a career that uses science, technology, engineering, and/or math (STEM) will allow me to get a job that is in high demand. |
| Interests | How much interest do you have in reading articles or books about engineering issues. | Rate your interest in reading about STEM issues. | Rate your interest in thinking about "hot topics" in any of the following: science, technology, engineering, and/or math (STEM). |

<sup>a</sup>(Lent et al., 2005)

**Table S2.**
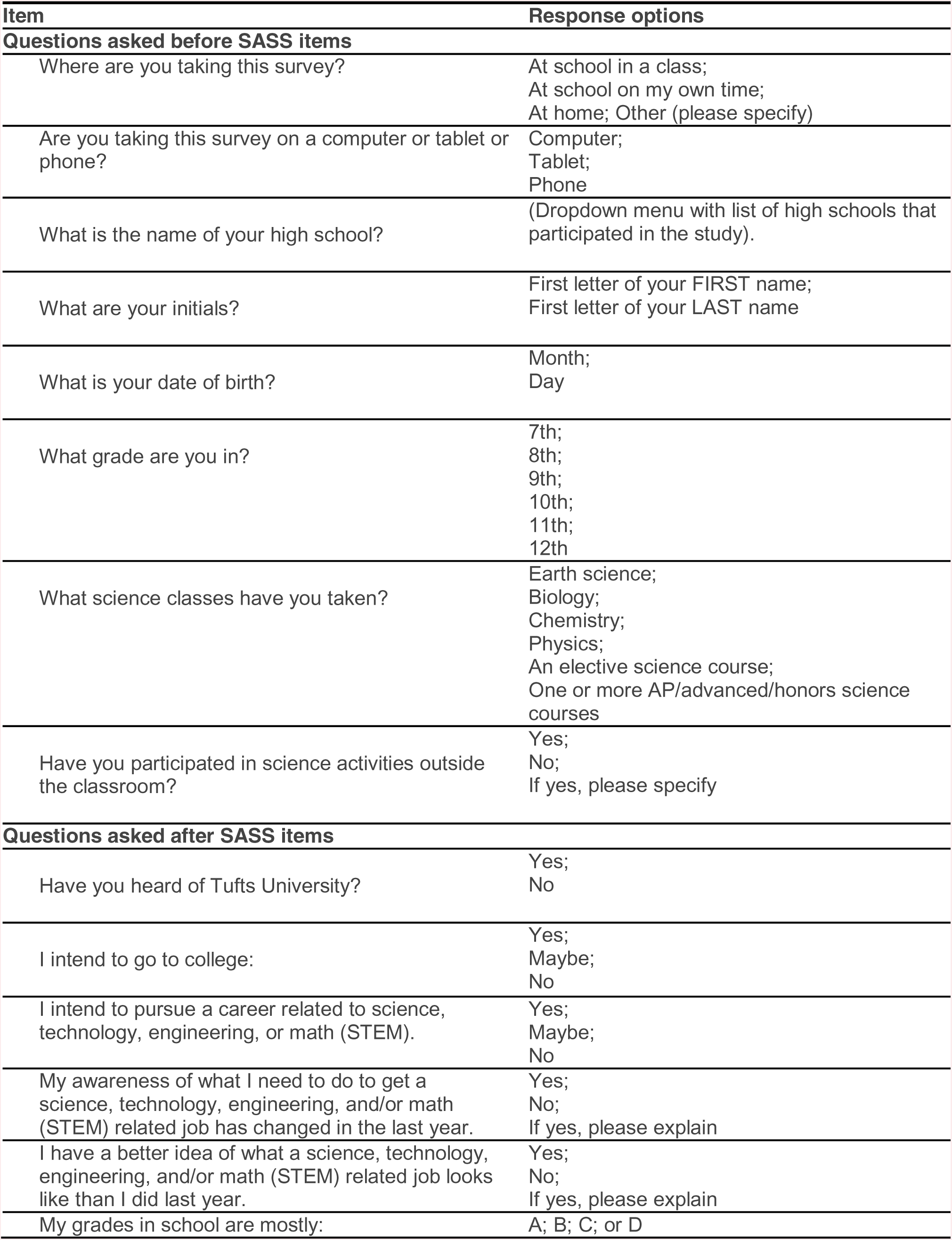
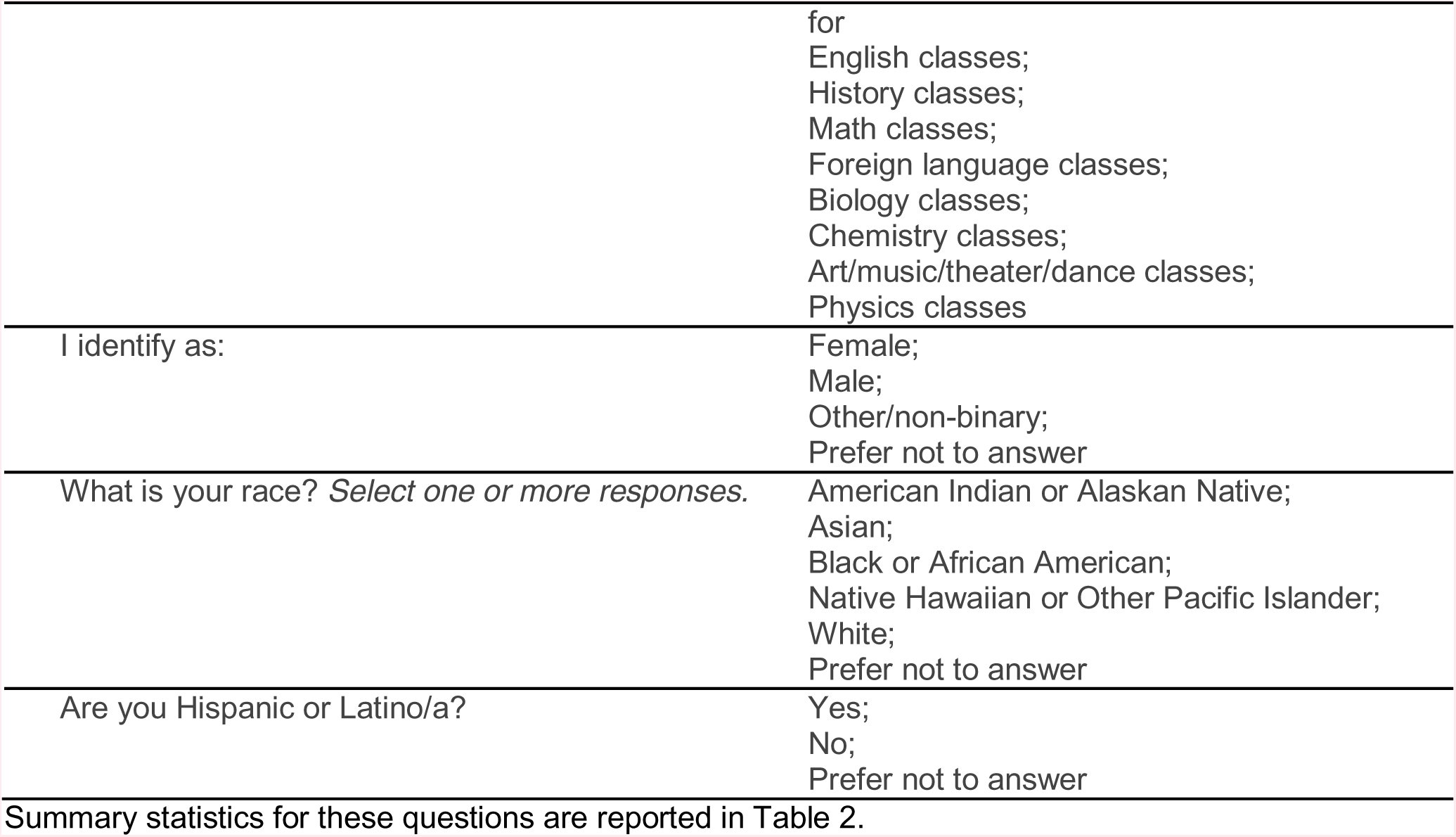
Background and demographic questions

**Table S3.** Item means and standard deviations for each sample

| Item | EFA Mean | EFA SD | CFA Mean | CFA SD |
| --- | --- | --- | --- | --- |
| Q3_1 | 4.42 | 1.13 | 4.38 | 1.10 |
| Q3_2 | 4.34 | 1.16 | 4.37 | 1.17 |
| Q3_3 | 4.01 | 1.27 | 4.02 | 1.27 |
| Q3_4 | 4.53 | 1.10 | 4.71 | 1.02 |
| Q3_5 | 4.51 | 1.06 | 4.62 | 1.12 |
| Q3_6 | 4.13 | 1.27 | 4.19 | 1.32 |
| Q9_1 | 4.99 | 1.11 | 4.96 | 1.19 |
| Q9_2 | 4.37 | 1.48 | 4.36 | 1.49 |
| Q9_3 | 4.25 | 1.52 | 4.41 | 1.50 |
| Q9_4 | 4.79 | 1.31 | 4.86 | 1.28 |
| Q10_1 | 4.24 | 1.30 | 4.32 | 1.26 |
| Q10_2 | 4.21 | 1.35 | 4.24 | 1.37 |
| Q12_1 | 4.97 | 1.05 | 5.13 | 1.01 |
| Q12_2 | 4.76 | 1.13 | 4.77 | 1.21 |
| Q12_3 | 4.79 | 1.17 | 4.83 | 1.24 |
| Q12_4 | 4.29 | 1.32 | 4.31 | 1.39 |
| Q12_5 | 4.82 | 1.24 | 4.87 | 1.23 |
| Q12_6 | 5.00 | 1.01 | 5.02 | 1.01 |
| Q13_1 | 4.66 | 1.33 | 4.78 | 1.22 |
| Q13_2 | 5.22 | 1.15 | 5.33 | 1.00 |
| Q13_3 | 5.14 | 1.14 | 5.25 | 0.99 |
| Q13_4 | 4.17 | 1.46 | 4.27 | 1.41 |
| Q14_1 | 4.64 | 1.22 | 4.51 | 1.37 |
| Q14_2 | 4.39 | 1.25 | 4.32 | 1.36 |
| Q14_3 | 4.17 | 1.33 | 4.19 | 1.39 |
| Q14_4 | 3.92 | 1.44 | 3.89 | 1.53 |
| Q14_5 | 4.20 | 1.36 | 4.10 | 1.39 |
| Q29_1 | 1.75 | 1.12 | 1.76 | 1.12 |
| Q29_2 | 2.21 | 1.36 | 2.28 | 1.46 |
| Q29_3 | 1.84 | 1.19 | 1.90 | 1.30 |
| Q16_1 | 4.24 | 1.29 | 4.21 | 1.33 |
| Q16_2 | 2.94 | 1.41 | 2.96 | 1.43 |
| Q16_3 | 2.36 | 1.28 | 2.28 | 1.23 |
| Q16_4 | 3.52 | 1.35 | 3.39 | 1.41 |
| Q16_5 | 3.15 | 1.51 | 3.35 | 1.54 |
| Q16_6 | 3.35 | 1.42 | 3.27 | 1.43 |
| Q16_7 | 2.88 | 1.36 | 2.82 | 1.35 |
| Q16_8 | 2.42 | 1.36 | 2.35 | 1.37 |
| Q16_9 | 3.67 | 1.50 | 3.65 | 1.48 |
| Q16_10 | 2.25 | 1.41 | 2.29 | 1.42 |
| SEEscore | NA | NA | 4.38 | 0.89 |
| SEAscore | NA | NA | 4.65 | 1.10 |
| OEscore | NA | NA | 4.86 | 0.84 |
| INscore | NA | NA | 4.20 | 1.13 |
| SPscore | NA | NA | 1.98 | 1.17 |
| CAscore | NA | NA | 3.06 | 0.93 |

Abbreviations: SD = standard deviation, SEE = Self-Efficacy-Experience, SEA = Self-Efficacy-Academic, OE = Outcome Expectations, IN = Interests, SP = Negative Perceptions of Scientists, CA = Career Awareness.

**Table S4.** Item loadings for five factor EFA

| Category | Item | PA4 | PA1 | PA5 | PA3 | PA2 | *h2* | *u2* | *com* |
| --- | --- | --- | --- | --- | --- | --- | --- | --- | --- |
| SE | Q3_1 | **0.42** | 0.00 | 0.25 | 0.05 | 0.09 | 0.33 | 0.67 | 1.77 |
| SE | Q3_2 | **0.61** | 0.00 | **0.32** | 0.07 | 0.02 | 0.60 | 0.40 | 1.56 |
| SE | Q3_3 | 0.05 | 0.04 | **0.72** | -0.08 | 0.02 | 0.61 | 0.39 | 1.04 |
| SE | Q3_4 | 0.17 | -0.01 | **0.59** | -0.01 | -0.04 | 0.42 | 0.58 | 1.18 |
| SE | Q3_5 | 0.10 | -0.06 | **0.56** | -0.04 | -0.01 | 0.33 | 0.67 | 1.10 |
| SE | Q3_6 | 0.08 | 0.01 | **0.79** | 0.02 | 0.03 | 0.69 | 0.31 | 1.02 |
| SE | Q9_1 | **0.45** | 0.15 | 0.25 | -0.07 | -0.04 | 0.46 | 0.54 | 1.92 |
| SE | Q9_2 | **0.58** | 0.06 | 0.11 | -0.11 | 0.02 | 0.47 | 0.53 | 1.17 |
| SE | Q9_3 | **0.70** | 0.05 | 0.03 | -0.01 | 0.00 | 0.53 | 0.47 | 1.02 |
| SE | Q9_4 | **0.76** | 0.07 | -0.16 | 0.00 | -0.02 | 0.57 | 0.43 | 1.11 |
| SE | Q10_1 | **0.32** | 0.11 | **0.41** | -0.07 | 0.14 | 0.54 | 0.46 | 2.40 |
| SE | Q10_2 | **0.65** | 0.02 | 0.05 | -0.01 | 0.10 | 0.50 | 0.50 | 1.06 |
| OE | Q12_1 | 0.14 | **0.58** | 0.05 | 0.09 | 0.01 | 0.44 | 0.56 | 1.18 |
| OE | Q12_2 | 0.00 | **0.66** | 0.09 | 0.14 | -0.01 | 0.46 | 0.54 | 1.13 |
| OE | Q12_3 | -0.05 | **0.48** | **0.35** | -0.05 | 0.01 | 0.49 | 0.51 | 1.89 |
| OE | Q12_4 | -0.07 | **0.55** | 0.24 | -0.01 | -0.03 | 0.43 | 0.57 | 1.41 |
| OE | Q12_5 | -0.03 | **0.77** | 0.06 | 0.07 | -0.03 | 0.58 | 0.42 | 1.03 |
| OE | Q12_6 | 0.01 | **0.73** | 0.05 | 0.08 | -0.04 | 0.54 | 0.46 | 1.04 |
| OE | Q13_1 | 0.05 | **0.58** | -0.02 | -0.11 | 0.13 | 0.45 | 0.55 | 1.19 |
| OE | Q13_2 | 0.08 | **0.75** | -0.14 | -0.10 | -0.03 | 0.58 | 0.42 | 1.14 |
| OE | Q13_3 | 0.08 | **0.77** | -0.17 | -0.11 | 0.03 | 0.62 | 0.38 | 1.16 |
| OE | Q13_4 | 0.11 | **0.52** | 0.17 | -0.04 | 0.07 | 0.50 | 0.50 | 1.38 |
| IN | Q14_1 | -0.04 | 0.27 | 0.27 | -0.11 | 0.12 | 0.29 | 0.71 | 2.78 |
| IN | Q14_2 | -0.08 | 0.23 | **0.40** | -0.13 | 0.18 | 0.42 | 0.58 | 2.42 |
| IN | Q14_3 | -0.10 | 0.26 | **0.42** | -0.10 | 0.29 | 0.55 | 0.45 | 2.79 |
| IN | Q14_4 | -0.02 | 0.11 | **0.43** | -0.02 | 0.21 | 0.37 | 0.63 | 1.61 |
| IN | Q14_5 | 0.08 | 0.19 | **0.40** | -0.08 | 0.23 | 0.48 | 0.52 | 2.27 |
| SP | Q29_1 | -0.06 | -0.01 | 0.05 | **0.86** | -0.02 | 0.75 | 0.25 | 1.02 |
| SP | Q29_2 | 0.03 | 0.06 | -0.05 | **0.80** | 0.03 | 0.63 | 0.37 | 1.02 |
| SP | Q29_3 | 0.03 | 0.00 | -0.03 | **0.90** | 0.03 | 0.80 | 0.20 | 1.01 |
| CA | Q16_1 | -0.07 | 0.18 | 0.17 | -0.03 | **0.41** | 0.35 | 0.65 | 1.84 |
| CA | Q16_2 | -0.14 | -0.06 | 0.05 | -0.03 | **0.55** | 0.30 | 0.70 | 1.18 |
| CA | Q16_3 | -0.10 | -0.02 | 0.01 | 0.04 | **0.67** | 0.43 | 0.57 | 1.05 |
| CA | Q16_4 | 0.08 | -0.04 | -0.03 | 0.07 | **0.54** | 0.29 | 0.71 | 1.11 |
| CA | Q16_5 | 0.14 | -0.03 | -0.09 | 0.06 | **0.64** | 0.40 | 0.60 | 1.17 |
| CA | Q16_6 | 0.02 | 0.01 | 0.06 | -0.04 | **0.74** | 0.61 | 0.39 | 1.02 |
| CA | Q16_7 | 0.10 | -0.02 | -0.09 | -0.01 | **0.84** | 0.68 | 0.32 | 1.06 |
| CA | Q16_8 | -0.03 | -0.01 | -0.03 | -0.03 | **0.68** | 0.44 | 0.56 | 1.01 |
| CA | Q16_9 | -0.04 | 0.11 | 0.11 | 0.04 | **0.60** | 0.46 | 0.54 | 1.16 |
| CA | Q16_10 | -0.07 | -0.05 | 0.06 | 0.04 | **0.66** | 0.44 | 0.56 | 1.06 |

**Table S5.** Item loadings for seven factor EFA

| Category | Item | PA5 | PA4 | PA1 | PA7 | PA6 | PA3 | PA2 | h2 | u2 | com |
| --- | --- | --- | --- | --- | --- | --- | --- | --- | --- | --- | --- |
| SEE | Q3_1 | **0.36** | **0.35** | -0.01 | 0.06 | -0.09 | 0.05 | 0.11 | 0.36 | 0.64 | 2.43 |
| SEE | Q3_2 | **0.43** | **0.54** | 0.10 | -0.05 | -0.15 | 0.04 | 0.07 | 0.65 | 0.35 | 2.24 |
| SEE | Q3_3 | **0.69** | -0.02 | 0.05 | 0.02 | 0.15 | -0.07 | 0.01 | 0.64 | 0.36 | 1.14 |
| SEE | Q3_4 | **0.70** | 0.06 | 0.04 | 0.00 | -0.06 | -0.03 | -0.03 | 0.52 | 0.48 | 1.05 |
| SEE | Q3_5 | **0.66** | -0.01 | 0.03 | -0.05 | -0.08 | -0.06 | 0.01 | 0.43 | 0.57 | 1.06 |
| SEE | Q3_6 | **0.76** | 0.00 | 0.09 | -0.04 | 0.13 | 0.02 | 0.02 | 0.73 | 0.27 | 1.09 |
| SEA | Q9_1 | 0.16 | **0.49** | 0.00 | 0.13 | 0.25 | -0.02 | -0.08 | 0.50 | 0.50 | 1.99 |
| SEA | Q9_2 | 0.01 | **0.65** | -0.04 | 0.04 | 0.21 | -0.07 | -0.02 | 0.53 | 0.47 | 1.25 |
| SEA | Q9_3 | -0.06 | **0.77** | -0.01 | 0.00 | 0.16 | 0.01 | -0.03 | 0.62 | 0.38 | 1.10 |
| SEA | Q9_4 | -0.04 | **0.74** | 0.03 | 0.06 | -0.15 | -0.02 | 0.01 | 0.55 | 0.45 | 1.11 |
| SEA | Q10_1 | **0.35** | **0.31** | 0.04 | 0.08 | 0.18 | -0.05 | 0.11 | 0.54 | 0.46 | 2.96 |
| SEA | Q10_2 | 0.13 | **0.62** | 0.03 | 0.01 | -0.09 | -0.02 | 0.12 | 0.50 | 0.50 | 1.22 |
| OE | Q12_1 | 0.08 | 0.14 | **0.50** | 0.16 | -0.05 | 0.04 | 0.04 | 0.46 | 0.54 | 1.50 |
| OE | Q12_2 | 0.01 | 0.02 | **0.69** | 0.03 | 0.03 | 0.08 | 0.02 | 0.52 | 0.48 | 1.04 |
| OE | Q12_3 | 0.17 | -0.02 | **0.55** | -0.06 | 0.21 | -0.09 | 0.01 | 0.54 | 0.46 | 1.60 |
| OE | Q12_4 | 0.08 | -0.04 | **0.66** | -0.08 | 0.14 | -0.07 | -0.02 | 0.52 | 0.48 | 1.19 |
| OE | Q12_5 | 0.04 | -0.02 | **0.72** | 0.15 | -0.01 | 0.00 | 0.00 | 0.62 | 0.38 | 1.10 |
| OE | Q12_6 | 0.07 | 0.01 | **0.67** | 0.17 | -0.05 | 0.02 | 0.01 | 0.58 | 0.42 | 1.17 |
| OE | Q13_1 | 0.00 | 0.09 | 0.24 | **0.42** | 0.13 | -0.08 | 0.10 | 0.49 | 0.51 | 2.18 |
| OE | Q13_2 | -0.07 | 0.12 | **0.30** | **0.60** | 0.08 | -0.07 | -0.04 | 0.68 | 0.32 | 1.67 |
| OE | Q13_3 | -0.02 | 0.09 | **0.33** | **0.64** | -0.05 | -0.09 | 0.04 | 0.75 | 0.25 | 1.63 |
| OE | Q13_4 | 0.13 | 0.13 | **0.43** | 0.15 | 0.09 | -0.06 | 0.08 | 0.50 | 0.50 | 1.90 |
| IN | Q14_1 | 0.09 | 0.04 | -0.11 | **0.36** | **0.56** | 0.00 | 0.00 | 0.52 | 0.48 | 1.86 |
| IN | Q14_2 | 0.14 | 0.00 | -0.03 | 0.20 | **0.62** | -0.04 | 0.05 | 0.59 | 0.41 | 1.35 |
| IN | Q14_3 | 0.09 | -0.01 | 0.15 | 0.02 | **0.60** | -0.04 | 0.19 | 0.67 | 0.33 | 1.40 |
| IN | Q14_4 | 0.06 | 0.08 | 0.17 | -0.20 | **0.59** | 0.02 | 0.11 | 0.55 | 0.45 | 1.55 |
| IN | Q14_5 | 0.02 | 0.19 | 0.23 | -0.17 | **0.57** | -0.05 | 0.14 | 0.66 | 0.34 | 1.93 |
| SP | Q29_1 | 0.05 | -0.06 | 0.03 | -0.04 | 0.00 | **0.86** | -0.02 | 0.75 | 0.25 | 1.02 |
| SP | Q29_2 | -0.08 | 0.05 | 0.06 | -0.01 | 0.04 | **0.81** | 0.01 | 0.65 | 0.35 | 1.04 |
| SP | Q29_3 | 0.00 | 0.03 | -0.03 | 0.03 | 0.00 | **0.92** | 0.01 | 0.85 | 0.15 | 1.01 |
| CA | Q16_1 | 0.24 | -0.10 | -0.01 | 0.27 | 0.08 | -0.01 | **0.40** | 0.40 | 0.60 | 2.72 |
| CA | Q16_2 | 0.07 | -0.16 | -0.06 | 0.01 | 0.01 | -0.03 | **0.54** | 0.30 | 0.70 | 1.26 |
| CA | Q16_3 | -0.05 | -0.09 | 0.11 | -0.14 | 0.03 | 0.01 | **0.67** | 0.46 | 0.54 | 1.20 |
| CA | Q16_4 | 0.03 | 0.06 | 0.00 | -0.02 | -0.07 | 0.05 | **0.55** | 0.30 | 0.70 | 1.08 |
| CA | Q16_5 | -0.07 | 0.13 | 0.10 | -0.12 | -0.10 | 0.01 | **0.67** | 0.45 | 0.55 | 1.27 |
| CA | Q16_6 | 0.05 | 0.01 | 0.01 | 0.01 | 0.08 | -0.05 | **0.72** | 0.61 | 0.39 | 1.04 |
| CA | Q16_7 | -0.09 | 0.11 | 0.02 | -0.03 | 0.01 | -0.03 | **0.83** | 0.69 | 0.31 | 1.06 |
| CA | Q16_8 | -0.04 | -0.03 | -0.06 | 0.06 | 0.07 | -0.02 | **0.65** | 0.44 | 0.56 | 1.07 |
| CA | Q16_9 | 0.17 | -0.06 | -0.07 | 0.24 | 0.08 | 0.07 | **0.57** | 0.51 | 0.49 | 1.68 |
| CA | Q16_10 | 0.12 | -0.10 | -0.12 | 0.10 | 0.02 | 0.05 | **0.65** | 0.46 | 0.54 | 1.25 |

**Table S6.** Item loadings for five factor CFA

| Item | SE | OE | IN | SP | CA |
| --- | --- | --- | --- | --- | --- |
| Q3_1 | 0.60 |  |  |  |  |
| Q3_2 | 0.73 |  |  |  |  |
| Q3_3 | 0.64 |  |  |  |  |
| Q3_4 | 0.62 |  |  |  |  |
| Q3_5 | 0.61 |  |  |  |  |
| Q3_6 | 0.68 |  |  |  |  |
| Q9_1 | 0.62 |  |  |  |  |
| Q9_2 | 0.67 |  |  |  |  |
| Q9_3 | 0.64 |  |  |  |  |
| Q9_4 | 0.52 |  |  |  |  |
| Q10_1 | 0.71 |  |  |  |  |
| Q10_2 | 0.62 |  |  |  |  |
| Q12_1 |  | 0.67 |  |  |  |
| Q12_2 |  | 0.65 |  |  |  |
| Q12_3 |  | 0.65 |  |  |  |
| Q12_4 |  | 0.69 |  |  |  |
| Q12_5 |  | 0.73 |  |  |  |
| Q12_6 |  | 0.71 |  |  |  |
| Q13_1 |  | 0.68 |  |  |  |
| Q13_2 |  | 0.66 |  |  |  |
| Q13_3 |  | 0.70 |  |  |  |
| Q13_4 |  | 0.61 |  |  |  |
| Q14_1 |  |  | 0.62 |  |  |
| Q14_2 |  |  | 0.77 |  |  |
| Q14_3 |  |  | 0.81 |  |  |
| Q14_4 |  |  | 0.75 |  |  |
| Q14_5 |  |  | 0.79 |  |  |
| Q29_1 |  |  |  | 0.87 |  |
| Q29_2 |  |  |  | 0.78 |  |
| Q29_3 |  |  |  | 0.90 |  |
| Q16_1 |  |  |  |  | 0.41 |
| Q16_2 |  |  |  |  | 0.47 |
| Q16_3 |  |  |  |  | 0.55 |
| Q16_4 |  |  |  |  | 0.54 |
| Q16_5 |  |  |  |  | 0.59 |
| Q16_6 |  |  |  |  | 0.82 |
| Q16_7 |  |  |  |  | 0.80 |
| Q16_8 |  |  |  |  | 0.68 |
| Q16_9 |  |  |  |  | 0.62 |
| Q16_10 |  |  |  |  | 0.63 |

**Table S7.**
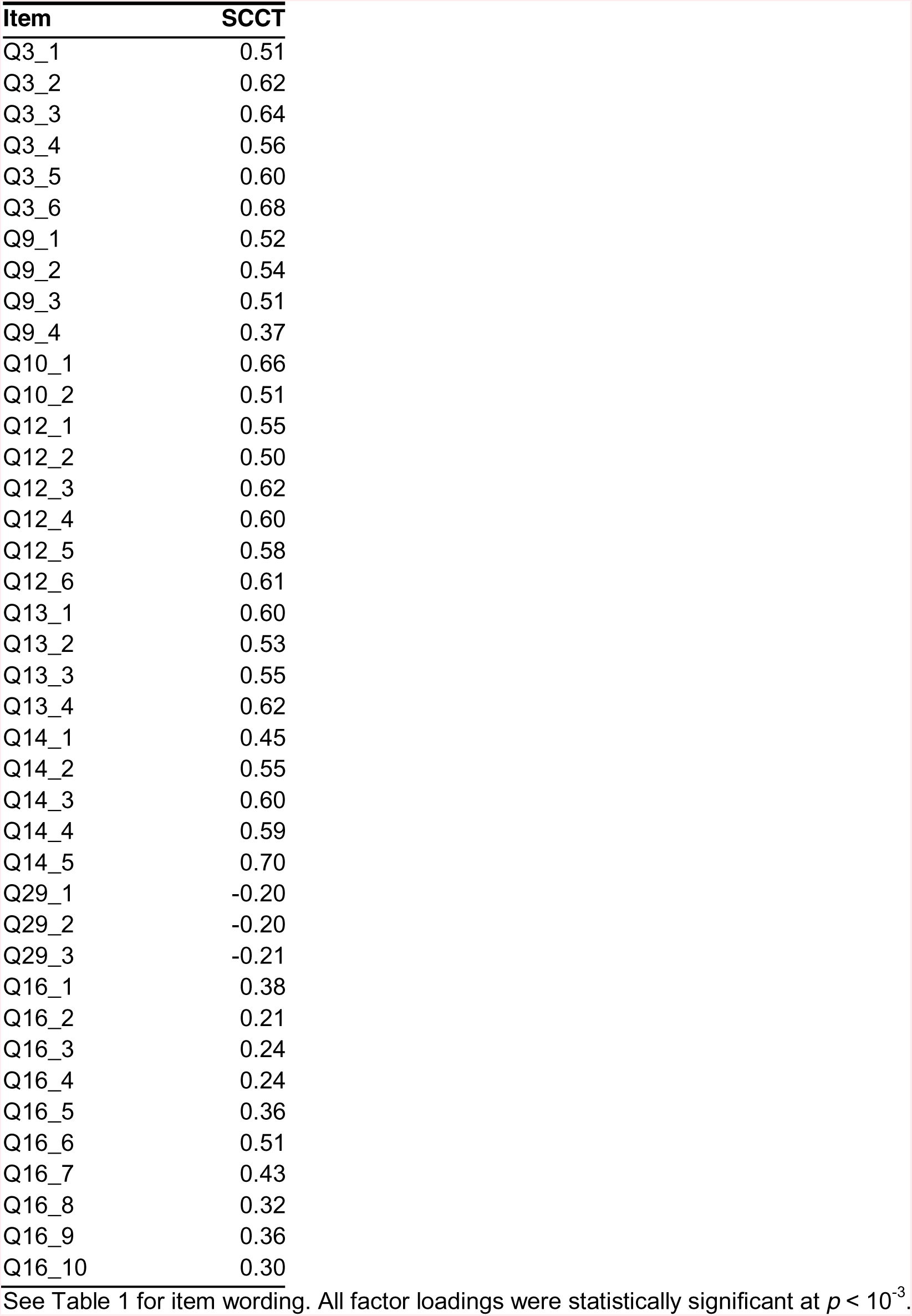
Item loadings for one factor CFA

| Item | SCCT |
| --- | --- |
| Q3_1 | 0.51 |
| Q3_2 | 0.62 |
| Q3_3 | 0.64 |
| Q3_4 | 0.56 |
| Q3_5 | 0.60 |
| Q3_6 | 0.68 |
| Q9_1 | 0.52 |
| Q9_2 | 0.54 |
| Q9_3 | 0.51 |
| Q9_4 | 0.37 |
| Q10_1 | 0.66 |
| Q10_2 | 0.51 |
| Q12_1 | 0.55 |
| Q12_2 | 0.50 |
| Q12_3 | 0.62 |
| Q12_4 | 0.60 |
| Q12_5 | 0.58 |
| Q12_6 | 0.61 |
| Q13_1 | 0.60 |
| Q13_2 | 0.53 |
| Q13_3 | 0.55 |
| Q13_4 | 0.62 |
| Q14_1 | 0.45 |
| Q14_2 | 0.55 |
| Q14_3 | 0.60 |
| Q14_4 | 0.59 |
| Q14_5 | 0.70 |
| Q29_1 | -0.20 |
| Q29_2 | -0.20 |
| Q29_3 | -0.21 |
| Q16_1 | 0.38 |
| Q16_2 | 0.21 |
| Q16_3 | 0.24 |
| Q16_4 | 0.24 |
| Q16_5 | 0.36 |
| Q16_6 | 0.51 |
| Q16_7 | 0.43 |
| Q16_8 | 0.32 |
| Q16_9 | 0.36 |
| Q16_10 | 0.30 |

See Table 1 for item wording. All factor loadings were statistically significant at $p < 10^{-3}$ .

**Table S8.** Dynamic fit CFA indices

| **Model** | **Level** | **CFI** | **RMSEA** | **SRMR** |
| --- | --- | --- | --- | --- |
| Five factor | Level 1: 95/5 | NONE | NONE | 0.036 |
|  | Level 1: 90/10 | NONE | NONE | -- |
|  | Level 2: 95/5 | 0.977 | 0.023 | 0.043 |
|  | Level 2: 90/10 | -- | -- | -- |
|  | Level 3: 95/5 | 0.948 | 0.035 | 0.053 |
|  | Level 3: 90/10 | -- | -- | -- |
|  | Level 4: 95/5 | 0.918 | 0.044 | 0.062 |
|  | Level 4: 90/10 | -- | -- | -- |
| Six Factor | Level 1: 95/5 | NONE | NONE | 0.035 |
|  | Level 1: 90/10 | NONE | NONE | -- |
|  | Level 2: 95/5 | 0.97 | 0.026 | 0.045 |
|  | Level 2: 90/10 | -- | -- | -- |
|  | Level 3: 95/5 | 0.945 | 0.036 | 0.054 |
|  | Level 3: 90/10 | -- | -- | -- |
|  | Level 4: 95/5 | 0.918 | 0.045 | 0.063 |
|  | Level 4: 90/10 | -- | -- | -- |
|  | Level 5: 95/5 | 0.889 | 0.053 | 0.071 |
|  | Level 5: 90/10 | -- | -- | -- |
| Reduced Six Factor | Level 1: 95/5 | NONE | NONE | 0.036 |
|  | Level 1: 90/10 | 0.991 | 0.015 | -- |
|  | Level 2: 95/5 | 0.974 | 0.026 | 0.045 |
|  | Level 2: 90/10 | -- | -- | -- |
|  | Level 3: 95/5 | 0.952 | 0.035 | 0.054 |
|  | Level 3: 90/10 | -- | -- | -- |
|  | Level 4: 95/5 | 0.924 | 0.045 | 0.063 |
|  | Level 4: 90/10 | -- | -- | -- |
|  | Level 5: 95/5 | 0.892 | 0.054 | 0.074 |
|  | Level 5: 90/10 | -- | -- | -- |

Abbreviations: *CFI* = comparative fit index, *RMSEA* = root-mean-square error of approximation, and *SRMR* = standardized root-mean-square residual.

**Table S9.** Item loadings for reduced six factor EFA and CFA

|  |  | EFA Results (*n* = 400) |  |  |  |  |  |  |  |  | CFA Results (*n* = 532) |  |  |  |  |  |
| --- | --- | --- | --- | --- | --- | --- | --- | --- | --- | --- | --- | --- | --- | --- | --- | --- |
| Category | Item | PA4 | PA5 | PA1 | PA6 | PA3 | PA2 | *h2* | *u2* | *com* | SEE | SEA | OE | IN | SP | CA |
| SEE | Q3_1 | **0.33** | **0.33** | 0.07 | -0.10 | 0.03 | 0.14 | 0.32 | 0.68 | 2.68 | 0.52 |  |  |  |  |  |
| SEE | Q3_3 | **0.62** | -0.01 | 0.06 | 0.21 | -0.08 | 0.01 | 0.61 | 0.39 | 1.28 | 0.77 |  |  |  |  |  |
| SEE | Q3_4 | **0.68** | 0.08 | 0.05 | -0.05 | -0.04 | 0.00 | 0.51 | 0.49 | 1.06 | 0.66 |  |  |  |  |  |
| SEE | Q3_5 | **0.70** | 0.04 | 0.00 | -0.10 | -0.07 | 0.04 | 0.48 | 0.52 | 1.07 | 0.69 |  |  |  |  |  |
| SEE | Q3_6 | **0.74** | 0.04 | 0.03 | 0.15 | 0.01 | 0.04 | 0.73 | 0.27 | 1.10 | 0.83 |  |  |  |  |  |
| SEA | Q9_1 | 0.14 | **0.51** | 0.09 | 0.21 | -0.02 | -0.07 | 0.50 | 0.50 | 1.65 |  | 0.73 |  |  |  |  |
| SEA | Q9_2 | 0.05 | **0.70** | -0.01 | 0.12 | -0.06 | 0.01 | 0.59 | 0.41 | 1.09 |  | 0.83 |  |  |  |  |
| SEA | Q9_3 | 0.01 | **0.85** | -0.02 | 0.05 | 0.04 | 0.00 | 0.73 | 0.27 | 1.01 |  | 0.80 |  |  |  |  |
| SEA | Q9_4 | 0.01 | **0.60** | 0.16 | -0.18 | -0.01 | 0.02 | 0.41 | 0.59 | 1.33 |  | 0.56 |  |  |  |  |
| OE | Q12_1 | 0.10 | 0.07 | **0.62** | -0.04 | 0.06 | 0.04 | 0.45 | 0.55 | 1.11 |  |  | 0.67 |  |  |  |
| OE | Q12_2 | 0.07 | -0.04 | **0.66** | 0.04 | 0.12 | 0.00 | 0.47 | 0.53 | 1.11 |  |  | 0.65 |  |  |  |
| OE | Q12_3 | 0.23 | -0.05 | **0.43** | 0.24 | -0.04 | -0.02 | 0.49 | 0.51 | 2.23 |  |  | 0.66 |  |  |  |
| OE | Q12_4 | 0.16 | -0.07 | **0.51** | 0.16 | -0.02 | -0.04 | 0.43 | 0.57 | 1.49 |  |  | 0.69 |  |  |  |
| OE | Q12_5 | 0.08 | -0.05 | **0.78** | -0.01 | 0.04 | -0.01 | 0.61 | 0.39 | 1.04 |  |  | 0.73 |  |  |  |
| OE | Q12_6 | 0.11 | -0.04 | **0.76** | -0.05 | 0.05 | 0.00 | 0.58 | 0.42 | 1.07 |  |  | 0.71 |  |  |  |
| OE | Q13_1 | -0.09 | 0.09 | **0.52** | 0.15 | -0.10 | 0.10 | 0.45 | 0.55 | 1.45 |  |  | 0.68 |  |  |  |
| OE | Q13_2 | -0.18 | 0.12 | **0.70** | 0.08 | -0.10 | -0.04 | 0.57 | 0.43 | 1.28 |  |  | 0.66 |  |  |  |
| OE | Q13_3 | -0.13 | 0.09 | **0.76** | -0.03 | -0.12 | 0.05 | 0.62 | 0.38 | 1.15 |  |  | 0.70 |  |  |  |
| OE | Q13_4 | 0.14 | 0.10 | **0.50** | 0.10 | -0.04 | 0.07 | 0.50 | 0.50 | 1.39 |  |  | 0.61 |  |  |  |
| IN | Q14_1 | -0.07 | 0.07 | 0.10 | **0.61** | -0.03 | -0.02 | 0.43 | 0.57 | 1.11 |  |  |  | 0.63 |  |  |
| IN | Q14_2 | 0.03 | 0.03 | 0.04 | **0.71** | -0.05 | 0.01 | 0.58 | 0.42 | 1.02 |  |  |  | 0.77 |  |  |
| IN | Q14_3 | 0.06 | 0.00 | 0.08 | **0.68** | -0.02 | 0.14 | 0.67 | 0.33 | 1.13 |  |  |  | 0.81 |  |  |
| IN | Q14_4 | 0.10 | 0.06 | -0.05 | **0.63** | 0.06 | 0.07 | 0.49 | 0.51 | 1.13 |  |  |  | 0.75 |  |  |
| IN | Q14_5 | 0.09 | 0.17 | 0.03 | **0.59** | 0.00 | 0.10 | 0.59 | 0.41 | 1.27 |  |  |  | 0.79 |  |  |
| SP | Q29_1 | 0.04 | -0.04 | -0.02 | 0.00 | **0.85** | -0.02 | 0.74 | 0.26 | 1.01 |  |  |  |  | 0.87 |  |
| SP | Q29_2 | -0.08 | 0.02 | 0.05 | 0.05 | **0.82** | 0.00 | 0.66 | 0.34 | 1.03 |  |  |  |  | 0.78 |  |
| SP | Q29_3 | -0.02 | 0.04 | -0.01 | -0.02 | **0.90** | 0.02 | 0.82 | 0.18 | 1.01 |  |  |  |  | 0.90 |  |
| CA | Q16_1 | 0.12 | -0.10 | 0.18 | 0.12 | -0.04 | **0.39** | 0.35 | 0.65 | 2.05 |  |  |  |  |  | 0.42 |
| CA | Q16_2 | 0.04 | -0.15 | -0.05 | 0.03 | -0.04 | **0.53** | 0.30 | 0.70 | 1.21 |  |  |  |  |  | 0.47 |
| CA | Q16_3 | -0.01 | -0.11 | -0.01 | 0.06 | 0.03 | **0.65** | 0.43 | 0.57 | 1.08 |  |  |  |  |  | 0.55 |
| CA | Q16_4 | 0.06 | 0.08 | -0.01 | -0.12 | 0.05 | **0.56** | 0.31 | 0.69 | 1.18 |  |  |  |  |  | 0.53 |
| CA | Q16_5 | 0.03 | 0.12 | 0.02 | -0.16 | 0.03 | **0.67** | 0.43 | 0.57 | 1.19 |  |  |  |  |  | 0.58 |
| CA | Q16_6 | 0.05 | 0.01 | 0.01 | 0.08 | -0.05 | **0.72** | 0.61 | 0.39 | 1.04 |  |  |  |  |  | 0.82 |
| CA | Q16_7 | -0.06 | 0.08 | 0.00 | 0.00 | -0.02 | **0.83** | 0.68 | 0.32 | 1.03 |  |  |  |  |  | 0.80 |
| CA | Q16_8 | -0.08 | -0.03 | -0.02 | 0.10 | -0.03 | **0.65** | 0.44 | 0.56 | 1.08 |  |  |  |  |  | 0.68 |
| CA | Q16_9 | 0.07 | -0.05 | 0.11 | 0.11 | 0.04 | **0.57** | 0.46 | 0.54 | 1.20 |  |  |  |  |  | 0.62 |
| CA | Q16_10 | 0.04 | -0.10 | -0.03 | 0.06 | 0.03 | **0.65** | 0.45 | 0.55 | 1.08 |  |  |  |  |  | 0.63 |

**Table S10.** CFA reduced six factor model latent variable correlations

|  | SEE | SEA | OE | IN | SP |
| --- | --- | --- | --- | --- | --- |
| SEA | 0.55* | - |  |  |  |
| OE | 0.58* | 0.44* | - |  |  |
| IN | 0.62* | 0.36* | 0.62* | - |  |
| SP | -0.17* | -0.19* | -0.20* | -0.16* | - |
| CA | 0.48* | 0.32* | 0.28* | 0.38* | -0.07 |

\* $p < 10^{-2}$ . Abbreviations: SEE = Self-Efficacy-Experience, SEA = Self-Efficacy-Academic, OE = Outcome Expectations, IN = Interests, SP = Negative Perceptions of Scientists, CA = Career Awareness.

**Table S11.** Measurement invariance testing results

| Group | Model | *robust* $\chi^2$ | $\chi^2$ | *df* | *CFI* | *TLI* | *RMSEA* | *SRMR* | $\Delta \chi^2$ | $\Delta df$ | *p-value* |
| --- | --- | --- | --- | --- | --- | --- | --- | --- | --- | --- | --- |
| Gender | Configural | 2705 | 3057 | 1228 | 0.82 | 0.81 | 0.07 | 0.07 |  |  |  |
|  | Metric | 2738 | 3098 | 1259 | 0.82 | 0.81 | 0.07 | 0.07 | 34 | 31 | 0.3054 |
|  | Scalar | 2973 | 3354 | 1290 | 0.78 | 0.79 | 0.08 | 0.07 | 258 | 31 | <2x10<sup>-16</sup> |
| Race | Configural | 2761 | 3035 | 1228 | 0.80 | 0.78 | 0.08 | 0.07 |  |  |  |
|  | Metric | 2790 | 3078 | 1259 | 0.80 | 0.79 | 0.08 | 0.08 | 34 | 31 | 0.3148 |
|  | Scalar | 2828 | 3113 | 1290 | 0.80 | 0.79 | 0.08 | 0.08 | 35 | 31 | 0.2873 |
|  | Conservative | 2910 | 3241 | 1327 | 0.79 | 0.79 | 0.08 | 0.08 | 82 | 37 | 2.9x10<sup>-5</sup> |
| SES | Configural | 2763 | 3034 | 1228 | 0.81 | 0.79 | 0.08 | 0.07 |  |  |  |
|  | Metric | 2787 | 3067 | 1259 | 0.81 | 0.80 | 0.08 | 0.07 | 27 | 31 | 0.6549 |
|  | Scalar | 2853 | 3133 | 1290 | 0.81 | 0.80 | 0.08 | 0.08 | 67 | 31 | 0.0002 |

For *CFI*, *TLI*, *RMSEA*, and *SRMR* robust measures were used. Abbreviations: *n* = number of students, $\chi^2$ = chi-square, *df* = degrees of freedom, *CFI* = comparative fit index, *TLI* = Tucker-Lewis index, and *RMSEA* = root-mean-square error of approximation, and *SRMR* = standardized root-mean-square residual. Both robust and standard chi-square values are reported; chi-square differences were based on the standard chi-square.

**Table S12.** Intraclass correlation coefficients

| Variable | *ICC(3,1)* | Lower Bound | Upper Bound |
| --- | --- | --- | --- |
| Q3 1 | 0.75 | 0.64 | 0.83 |
| Q3 2 | 0.75 | 0.64 | 0.83 |
| Q3 3 | 0.60 | 0.44 | 0.72 |
| Q3 4 | 0.59 | 0.43 | 0.71 |
| Q3 5 | 0.37 | 0.18 | 0.54 |
| Q3 6 | 0.48 | 0.30 | 0.63 |
| Q9 1 | 0.69 | 0.56 | 0.78 |
| Q9 2 | 0.90 | 0.85 | 0.93 |
| Q9 3 | 0.95 | 0.92 | 0.96 |
| Q9 4 | 0.60 | 0.44 | 0.72 |
| Q10 1 | 0.51 | 0.34 | 0.65 |
| Q10 2 | 0.61 | 0.46 | 0.73 |
| Q12 1 | 0.44 | 0.25 | 0.59 |
| Q12 2 | 0.53 | 0.36 | 0.66 |
| Q12 3 | 0.39 | 0.20 | 0.56 |
| Q12 4 | 0.43 | 0.24 | 0.59 |
| Q12 5 | 0.50 | 0.32 | 0.64 |
| Q12 6 | 0.52 | 0.35 | 0.66 |
| Q13 1 | 0.45 | 0.26 | 0.60 |
| Q13 2 | 0.63 | 0.48 | 0.74 |
| Q13 3 | 0.44 | 0.25 | 0.59 |
| Q13 4 | 0.58 | 0.42 | 0.70 |
| Q14 1 | 0.63 | 0.49 | 0.74 |
| Q14 2 | 0.54 | 0.38 | 0.68 |
| Q14 3 | 0.52 | 0.35 | 0.66 |
| Q14 4 | 0.50 | 0.33 | 0.65 |
| Q14 5 | 0.56 | 0.40 | 0.69 |
| Q29 1 | 0.64 | 0.50 | 0.75 |
| Q29 2 | 0.68 | 0.55 | 0.78 |
| Q29 3 | 0.63 | 0.49 | 0.75 |
| Q16 1 | 0.61 | 0.46 | 0.73 |
| Q16 2 | 0.60 | 0.45 | 0.72 |
| Q16 3 | 0.42 | 0.23 | 0.58 |
| Q16 4 | 0.61 | 0.46 | 0.73 |
| Q16 5 | 0.78 | 0.68 | 0.85 |
| Q16 6 | 0.41 | 0.23 | 0.57 |
| Q16 7 | 0.43 | 0.24 | 0.59 |
| Q16 8 | 0.30 | 0.10 | 0.48 |
| Q16 9 | 0.48 | 0.30 | 0.63 |
| Q16 10 | 0.25 | 0.04 | 0.43 |
| SEE score | 0.64 | 0.50 | 0.75 |
| SEA score | 0.90 | 0.85 | 0.93 |
| OE score | 0.53 | 0.36 | 0.67 |
| IN score | 0.57 | 0.41 | 0.70 |
| SP score | 0.72 | 0.60 | 0.81 |
| CA score | 0.47 | 0.29 | 0.62 |

## Supplemental Figures

**Figure S1.**
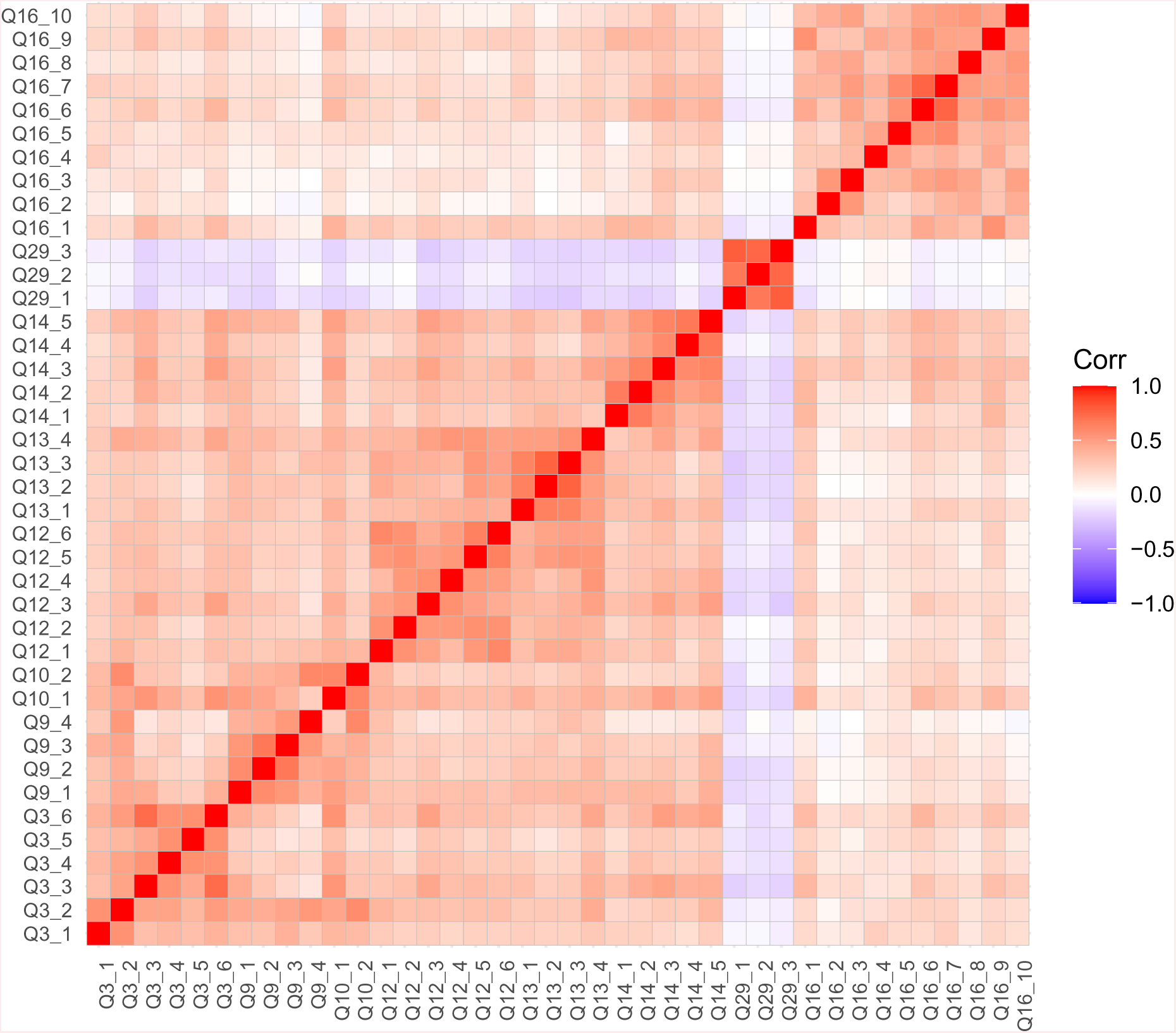
EFA correlation plot An inter-item correlation plot was used to support factorability of the dataset. Items with positive correlations are shaded in red, while those with negative correlations are blue. Intensity of the color indicates the strength of the correlation. At least six factors can be identified. Several items exhibit correlations between 0.2 to 0.8. For item wording, see Table 1. *n* = 400.

**Figure S2.**
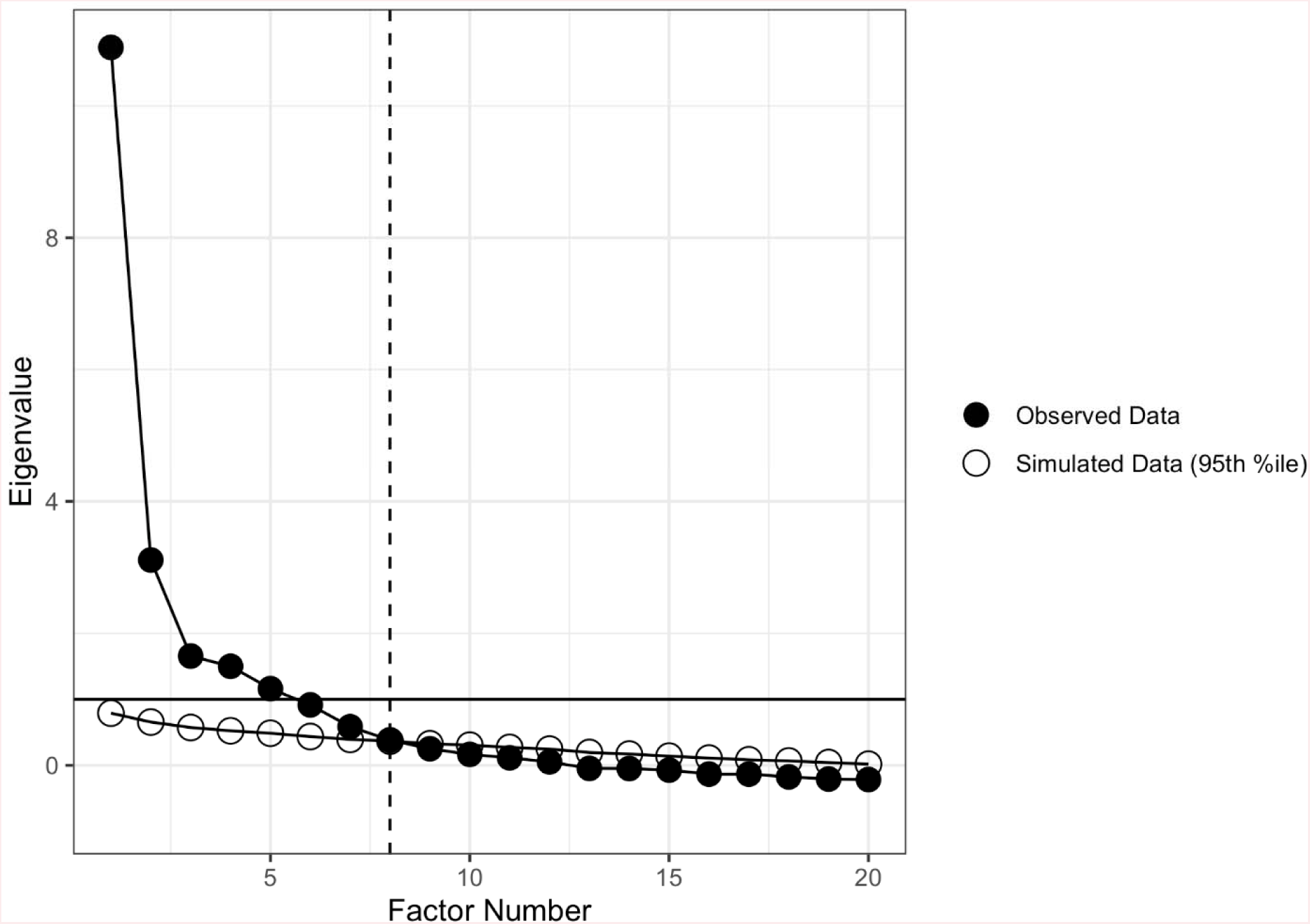
Scree plot Scree plot of eigenvalues (variance explained; descending order) for each factor that could be extracted from the data (filled circles). Eigenvalues for factors extracted from parallel analysis are also shown (open circles).

**Figure S3.**
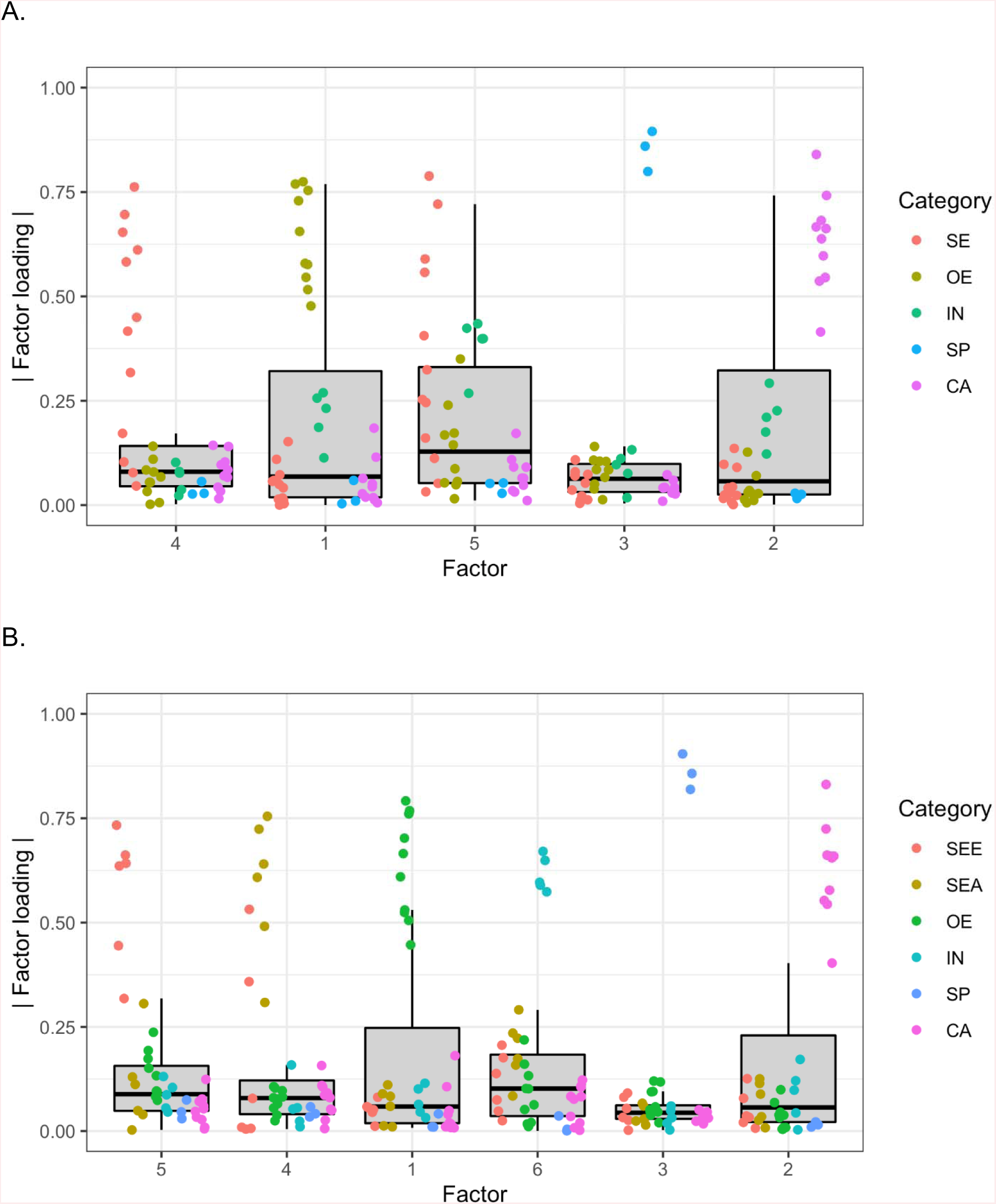
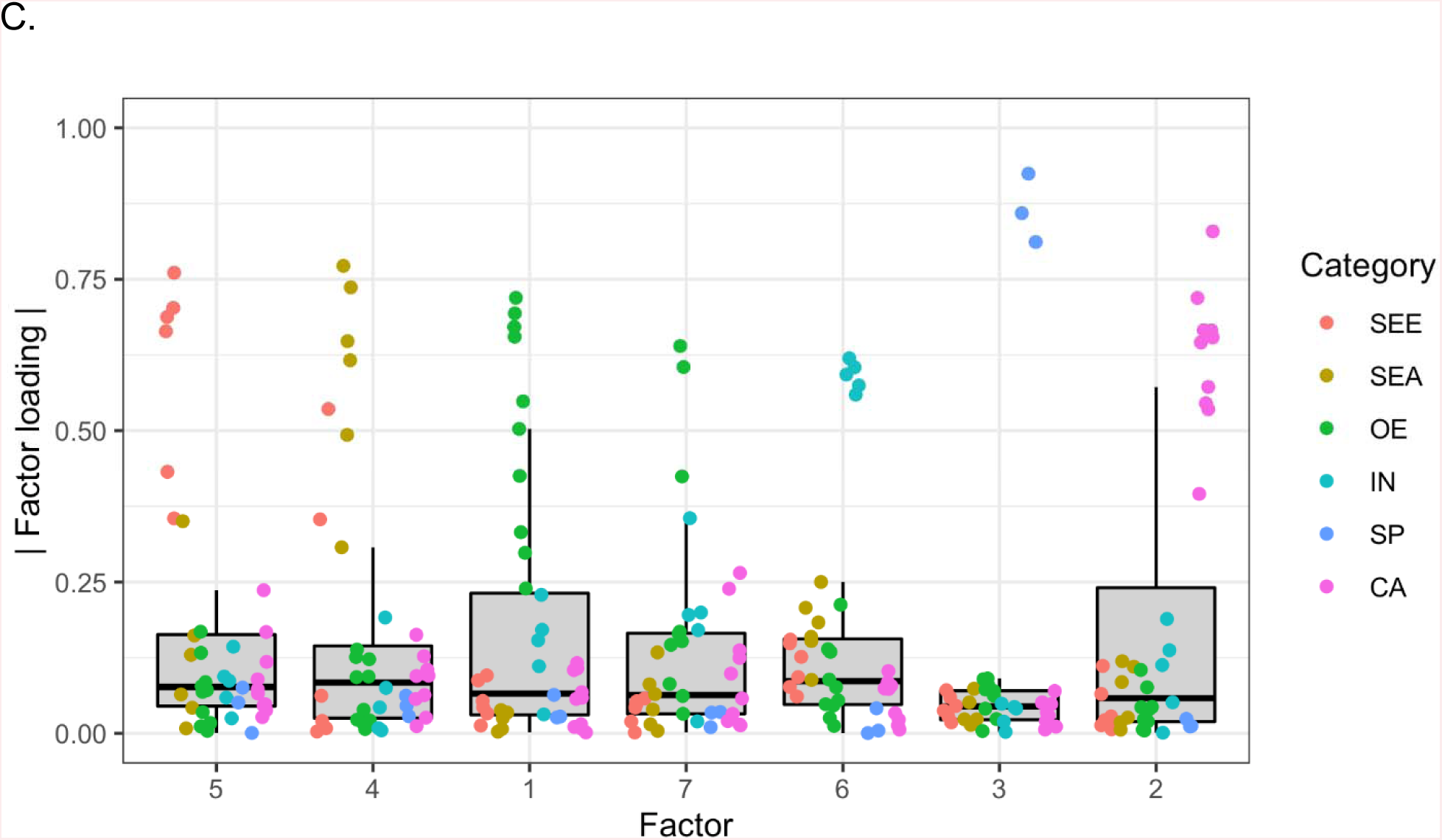
EFA box plots EFA results were visualized using box plots for the A) five factor, B) six factor models, and C) seven factor models. The absolute value of each item’s factor loading was plotted for each factor. Items were colored according to their a priori determined factor. Abbreviations: SE = Self-Efficacy, SEE = Self-Efficacy-Experience, SEA = Self-Efficacy-Academic, OE = Outcome Expectations, IN = Interests, SP = Negative Perceptions of Scientists, CA = Career Awareness.

**Figure S4.**
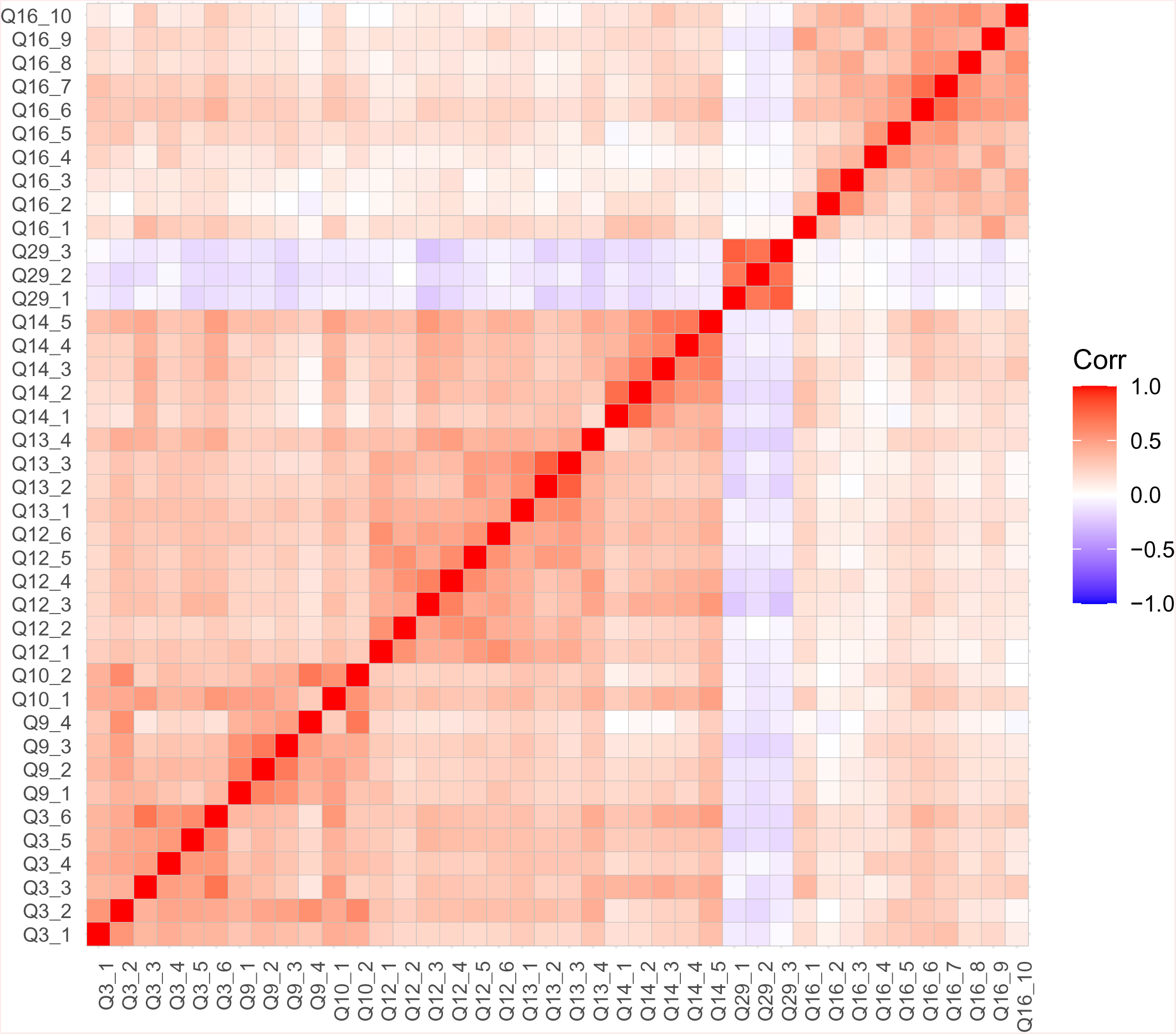
CFA correlation plot An inter-item correlation plot was used to support factorability of the dataset. Items with positive correlations are shaded in red, while those with negative correlations are blue. Intensity of the color indicates the strength of the correlation. At least six factors can be identified. Several items exhibit correlations between 0.2 to 0.8. See Table 1 for item wording. *n* = 532.

**Figure S5.**
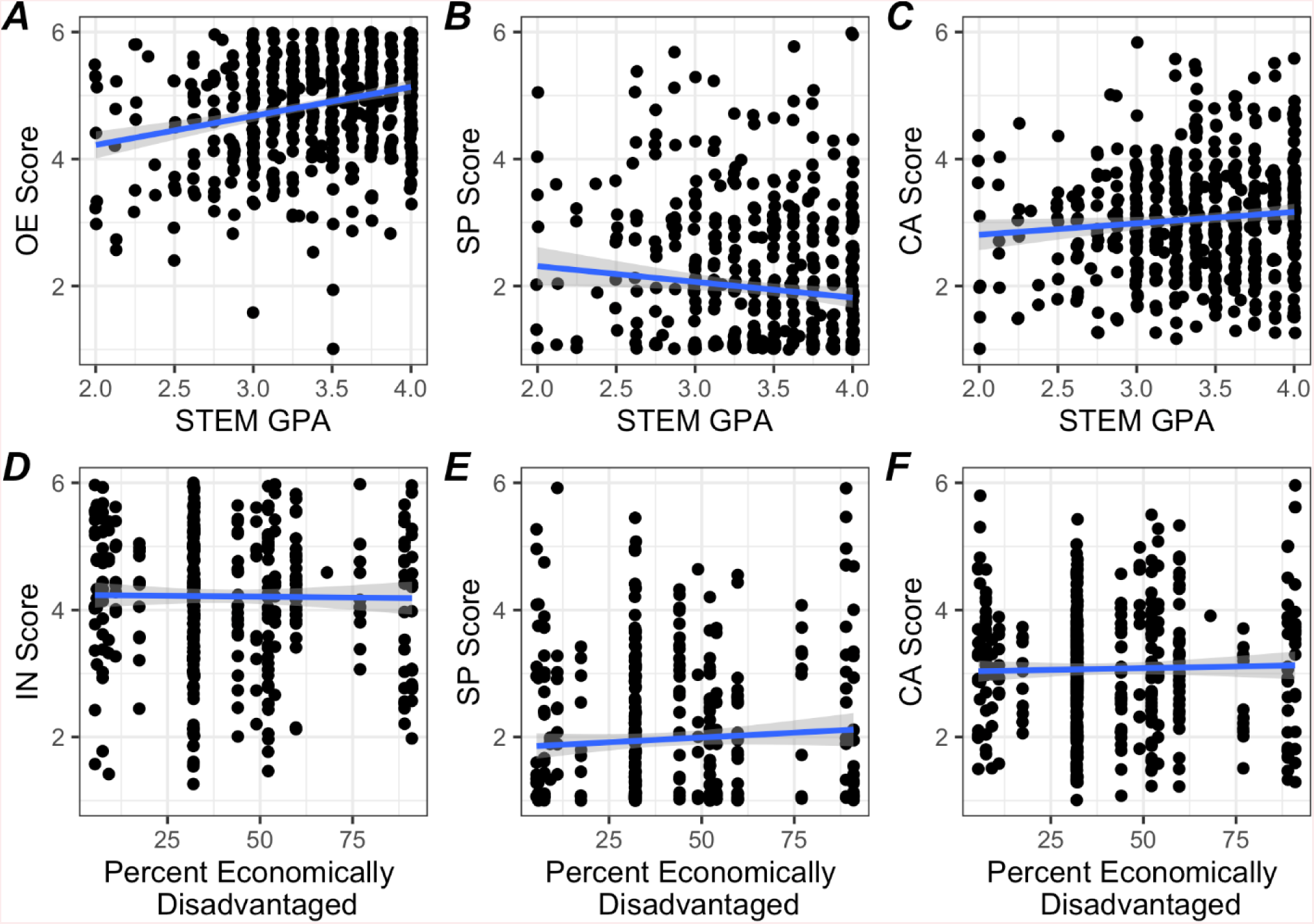
Additional criterion-related validity testing Scatter plots illustrating the relationship of factor scores and external variables are shown with linear regression lines. Spearman’s correlations and *p* values were also calculated for SASS factors with STEM GPA: A) Outcome Expectations score (*rho* = 0.24, *p* < 10^-7^); B) Negative Perceptions of Scientists score (*rho* = -0.10, *p* = 0.03); C) Career Awareness score (*rho* = 0.09, *p* = 0.03); and for SASS factors with school percent economically disadvantaged: D) Interests score (*rho* = -0.01, *p* = 0.8); E) Negative Perceptions of Scientists score (*rho* = 0.06, *p* = 0.2); and F) Career Awareness score (*rho* = 0.02, *p* = 0.6). A high percentage of economically disadvantaged students corresponds with low SES for the school. Because the SES data is collected at the school level, the values are clustered at discrete values on the horizontal axis. Abbreviations: SEE = Self-Efficacy-Experience, SEA = Self-Efficacy-Academic, OE = Outcome Expectations, IN = Interests, SP = Negative Perceptions of Scientists, CA = Career Awareness.

**Figure S6.**
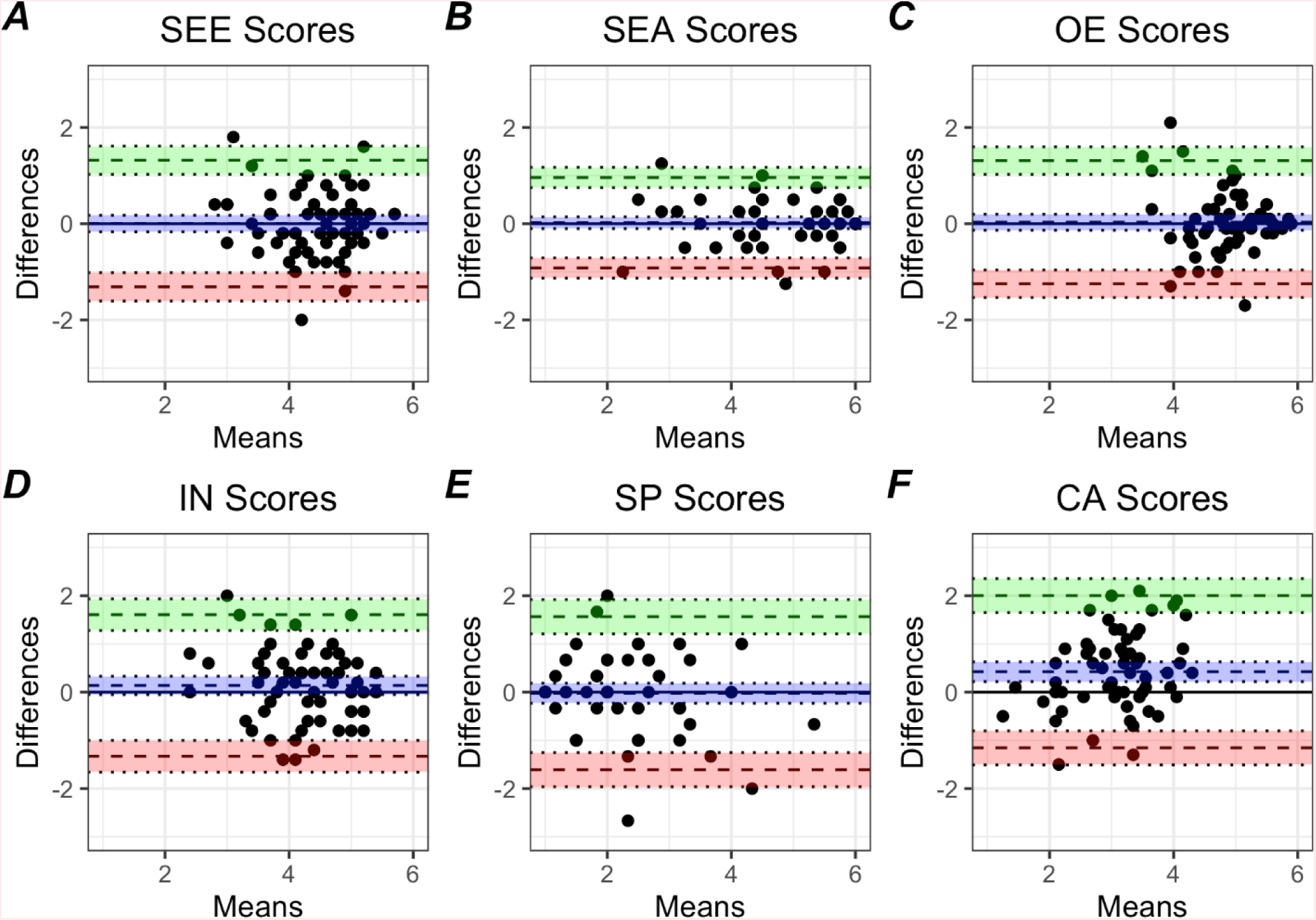
Bland-Altman plots Bland-Altman plots were generated to evaluate the agreement between student factor scores at each timepoint. For each factor, the differences between each score at each timepoint was calculated and plotted against the mean for each timepoint. The biases, or the average of the distances between the measurements (A: 0.008, B: 0.02, C: 0.03, D: 0.14, E: -0.02, and F: 0.42) and the lower and upper limits of agreement (A: -1.17 and 1.19, B: -0.91 and 0.96, C: -1.25 and 1.31, D: -1.34 and 1.61, E: -1.61 and 1.57, and F: -1.15 and 2.00) are represented as dashed lines. The shaded areas represent 95% confidence intervals for the biases (purple), upper limits of agreement (green), and lower limits of agreement (red). Abbreviations: SEE = Self-Efficacy-Experience, SEA = Self-Efficacy-Academic, OE = Outcome Expectations, IN = Interests, SP = Negative Perceptions of Scientists, CA = Career Awareness.

